# Linking neuron-axon-synapse architecture to white matter vasculature using high-resolution multimodal MRI in primate brain

**DOI:** 10.1101/2025.04.15.648990

**Authors:** Ikko Kimura, Takuya Hayashi, Joonas A. Autio

## Abstract

Blood vessels and axons align outside the brain due to shared growth factors. However, this neuron-axon-synapse and vessel relationship within the brain white matter remains unclear, primarily due to the technical challenges of charting the complex trajectories of fiber tracts and the dense network of arteries. Consequently, the organizational logic and neurometabolic factors shaping white matter vasculature remain poorly understood. Here, we address these questions using high-resolution multimodal MRI, in vitro neuron density, and receptor autoradiography in macaque monkeys. In superficial white matter, vascularity exhibited parallel alignment with the cortical surface. This vascularity showed negligible dependence on overlying gray matter neuron density (R^2^ = 0.01), minimal dependence on white matter myelination (R^2^ = 0.10), and moderate correlation with receptor density (R^2^ = 0.27). These suggest an association of vascularity with energy demands and axonal branching. In deep white matter, axon geometry, density, and proximity to the cortical surface predict vascular volume with high precision (R^2^ = 0.62). Overall, these findings establish a relation between neuron-axon-synapse architecture and white matter vasculature in the primate brain, offering advances in understanding the organization and pathophysiology of white matter.

## 1. Introduction

White matter vasculature plays a crucial role in brain function and aging-related diseases. Nevertheless, its organization and function remain underexplored, despite supplying a substantial proportion of brain’s blood flow (Wu et al., 2013). A deeper understanding of white matter vascularity is essential for elucidating its contributions to neural function (Autio and Roberts, 2014; Huang et al., 2023, 2020; Schilling et al., 2023; Wang et al., 2023; Weber, 2002), plasticity (Kimura et al., 2024; Sampaio-Baptista and Johansen-Berg, 2017), and pathology (Chen et al., 2013; Chojdak-Łukasiewicz et al., 2021). Early post-mortem studies mapped the arterial supply of white matter from pial vessels (Duvernoy et al., 1981), and research outside the brain showed that blood vessels and axons align due to shared growth factors (Carmeliet and Tessier-Lavigne, 2005). However, the fundamental principles governing the brain white matter vascular organization remain poorly understood (Smirnov et al., 2021), limiting insights into its role in brain health and disease.

The neurometabolic factors influencing white matter vasculature also remain uncertain. Energy demand, reflected by mitochondria density, varies among tracts (Kageyama and Wong-Riley, 1986, 1984; Perge et al., 2012), and regions with high energy demands tend to have denser vascularization (Borowsky and Collins, 1989; Perge et al., 2012). Energy consumption in white matter stems from several biophysical processes, including protein and lipid synthesis, axonal transport of molecules and organelles (Bernstein and Bamburg, 2003) and, to a lesser extent, axonal action potentials (Harris and Attwell, 2012). Although myelin is generally thought to conserve energy through saltatory conduction, simulation studies suggest that the energetic costs of myelin synthesis and oligodendrocyte maintenance may offset these savings (Harris and Attwell, 2012). Variations in gray matter neural and receptor density may further impact white matter energy demands, as axonal transport is essential for moving synaptic vesicles between the soma and axon terminals (Bernstein and Bamburg, 2003; Goldstein et al., 2008). However, empirical evidence linking neurometabolic factors to white matter vascularization remains limited.

Advancements in high-resolution ferumoxytol-weighted MRI have significantly improved the quantification of brain vascularity (Autio et al., 2025; Bernier et al., 2019). Our recent work demonstrated that cortical layer vasculature was related to regional variations in neuron, and synaptic or myelin densities (Collins et al., 2010; Froudist-Walsh et al., 2023) in macaque monkeys (Autio et al., 2025). Additionally, diffusion MRI (dMRI) provides crucial insights into the organization and microstructure of white matter tracts, including neurite density, axon orientation and its dispersion (Basser et al., 1994; Zhang et al., 2012), enabling direct comparisons between fiber architecture and vascular organization.

In this study, we applied multimodal MRI to investigate the neurometabolic factors influencing white matter vascular organization in macaques. By estimating vascular volume in both superficial and deep white matter, we identified substantial inter-areal variability, comparable to that seen in cortical gray matter. To understand the anatomical basis of this variability, we examined the relationship between white matter vasculature and microstructure, as well as the neuronal and synaptic architecture of overlying cortical areas. As a preview, we found that: 1) white matter regions with higher myelin exhibited lower vascular density, suggesting reduced energy demands, though this explains only a small portion of the variability, consistent with the result of a simulation study (Harris and Attwell, 2012); 2) white matter vascularity correlated more strongly with overlying cortical receptor density than with neuron density; and 3) the geometrical arrangement, density, and distance of axons to the cortical surface predict white matter vasculature with high precision. These findings provide new insights into the metabolic and structural determinants of white matter vascular organization.

## 2. Methods

### 2.1 Animals

MRI data from four macaques (Macaca mulatta, age: 4 – 6 years, weight: 7.4 – 8.4 kg) (Autio et al., 2025) were used to investigate the vascularity and microstructure of the white matter in the brain. All experiments were approved by the Animal Care and Use Committee of the Kobe Institute of RIKEN (MA2008-03-11) and conducted in accordance with the institutional guidelines for animal experiments, the Basic Policies for the Conduct of Animals Experiments in Research Institution (MEXT, Tokyo, Japan), and the Guidelines for the Care and Use of Laboratory Animals (ILAR, Washington DC, USA).

### 2.2 Image acquisition

Structural, ferumoxytol-weighted multi-echo gradient-echo and dMRI data were sourced from our prior studies (Autio et al., 2025, 2024, 2020) using the same macaques across modalities (n = 4). All MRI scans were acquired using a Siemens Prisma 3T scanner (Siemens, Erlangen, Germany) equipped with a 24-channel multi-array RF coil for non-human primate brains (Rogue Research, Montreal, Canada / Takashima Seisakusho KK, Tokyo, Japan) (Autio et al., 2020). The animals were scanned under anesthesia and physiological conditions were continuously monitored throughout the experiments. Detailed experimental procedures and imaging protocols can be found in (Autio et al., 2024) for structural MRI, in (Autio et al., 2025) for multi-echo gradient-echo MRI, and in (Autio et al., 2020) for dMRI.

In brief, multi-echo gradient-echo MRI data with 10 equidistant echo-times (TE = 3.4 – 25.0 ms, TE interval = 2.4 ms) were acquired before (2 scans per trial) and after (5 scans per trial) an intravascular injection of 12 mg/kg ferumoxytol to investigate brain vascularity (0.32 mm isotropic resolution). A subset of these macaques was rescanned after ferumoxytol injection using three equidistant TEs (TE = 5.0 – 13.0 ms, interval = 4.0 ms) with spatial resolution adjusted to match the critical (spatial) sampling frequency of intra-cortical vessels (0.23 mm isotropic resolution) (Autio et al., 2025; Weber et al., 2008; Zheng et al., 1991). Structural T1-weighted (T1w) MPRAGE and T2-weighted (T2w) SPACE-FLAIR images were acquired at 0.32 mm isotropic resolution to accurately reconstruct white matter and pial surfaces. Additionally, T1w/T2w-FLAIR was used as an indirect proxy measure to investigate myeloarchitecture across cortical layers and within the adjacent superficial white matter, suppressing B_1_ receive bias (Glasser et al., 2013) and reducing Gibbs ringing artifacts when using FLAIR-modified sequences (Autio et al., 2024). The Human Connectome Project (HCP)-style diffusion MRI data acquisition protocol was applied to study white matter tract orientation and microstructure (Autio et al., 2020; Glasser et al., 2013). These images were obtained at 0.9 mm isotropic resolution with b-values of 0, 1000, 2000, and 3000 mm/s^2^, using a total of 500 non-collinear diffusion orientations. These directions were uniformly distributed over the whole sphere in each non-zero b-value shell.

### 2.3 Image Analysis

Structural and dMRI data were preprocessed using a non-human primate version of the preprocessing pipelines developed by the HCP (https://github.com/Washington-University/NHPPipelines) (Autio et al., 2020; Donahue et al., 2016; Glasser et al., 2013; Hayashi et al., 2021). These pipelines utilize FreeSurfer (version 6.0.1), wb_command (version 1.5.0), and the FMRIB software library (FSL; version 6.0.4). Detailed preprocessing procedures for structural and diffusion MRI can be found in (Autio et al., 2020).

Briefly, T1w and T2w-FLAIR images were aligned to an anterior-posterior commissural (AC-PC) space, corrected for B_1_-bias field, and nonlinearly registered to the Mac30BS standard space (Hayashi et al., 2021). The pial and white surfaces and subcortical structures were delineated using FreeSurfer. These surfaces were registered to the Mac30BS standard space based on sulcus landmarks using MSMSulc, and a mid-thickness surface was created. All surfaces were visually inspected for errors. When required, manual corrections were made on a wm.edit.mgz or subtle manual repositioning was performed to a surface using FreeSurfer 7.1 (by JAA).

Using the corrected white matter and pial surfaces as references, we generated a layer for superficial white matter positioned 0.32 mm beneath the white matter surface (Kirilina et al., 2020). We additionally created 12 equivolumetric layers (ELs) (Autio et al., 2025, 2024; Bok, 1929; Van Essen and Maunsell, 1980). These ELs were positioned within the cerebral cortex (i.e., EL1a, EL1b, EL2a, … EL6a, and EL6b), with EL1a situated just beneath the pial surface and EL6b situated just above the white matter surface. This nomenclature was intended to facilitate comparison with anatomical cortical layers (e.g., Ia, Ib, Ic, … VIb).

Vascular volume was evaluated using ferumoxytol-weighted multi-echo gradient-echo imaging (Autio et al., 2025). Quantitative R_2_* fitting was performed on the multi-echo gradient-echo images before and after ferumoxytol injection using hMRI toolbox (Tabelow et al., 2019). The pre-ferumoxytol R_2_* maps were subtracted from post-ferumoxytol maps to generate ΔR_2_* maps, providing indirect measures of vascular volume (Boxerman et al., 1995; Kim et al., 2013). These ΔR_2_* volume maps were then mapped onto native cortical layers using ribbon-constrained volume-to-surface-mapping in wb_command and resampled to 32k and 164k meshes. Subsequently, these ΔR_2_* maps were corrected for the orientation of the cerebral cortex relative to the direction of the static magnetic field (B_0_) (Autio et al., 2025). The median value of each area in the M132 Lyon macaque brain atlas (Markov et al., 2014) or Julich macaque brain atlas (Froudist-Walsh et al., 2023) was calculated and averaged across the macaques (n = 4). The M132 atlas was used to investigate whole brain relationships between vasculature and myelin index whereas the Julich atlas was additionally used to align with publicly available datasets on neuron and receptor densities in the overlying gray matter (See *2.4 Statistical Analysis* for details). In the M132 Lyon atlas (n = 91 per hemisphere), areas 24a, 24b, 29_30, and V2 were excluded from further analysis (resulting in n = 87 per hemisphere), and, in the Julich atlas (n = 109, left hemisphere), areas a24ab, p24ab, 24ab, 29/30, V2v, and V2d were excluded (resulting in n = 103) due to minor yet significant errors in white surface estimation. To visualize white matter macrovessels, post-ferumoxytol images with high spatial resolution (0.23 mm isotropic) were averaged across runs and TEs to improve signal-to-noise ratio. A Hessian-based Frangi ‘vesselness’ filter (https://jp.mathworks.com/matlabcentral/fileexchange/24409-hessian-based-frangi-vesselness-filter) was then applied to detect vessels in the 3D volumetric images. To enable surface visualization of vessels, the 164k surface mesh was resampled to ultra-high resolution 656k mesh, resulting in an average 0.14 mm spacing between vertices in the superficial white matter. In cortical gray matter, this spacing is smaller than the average distance between the penetrating vessels (Autio et al., 2025; Weber et al., 2008). To visualize the macrovessels, spatial gradients on each layer were calculated using wb_command -metric-gradient.

dMRI data were intensity-normalized, distortion corrected, and corrected for eddy current and subject motion. The corrected images were co-registered to the native AC-PC aligned T1w image. To investigate the relationship between vascularity and major fiber bundles, tractography was performed to define the location of commissural fibers connecting between bilateral identical areas. We selected these bundles for analysis because they can be robustly delineated and consistently traverse from the cerebral cortex to the deep white matter. Tractography was performed using MRtrix3 (version 3.0.4) (Tournier et al., 2019). First, the basis functions to deconvolve dMRI data to create the fiber orientation density (FOD) were calculated with the dhollander algorithm (i.e., separate basis functions were estimated for gray matter, white matter, and cerebrospinal fluid and for each b value). These basis functions were used to calculate an FOD on each voxel with multi-shell multi-tissue constrained spherical deconvolution (MSMT-CSD). The tissue boundary between the white and cortical gray matter for seeding tractography was created with preprocessed structural MRI data. One hundred million streamlines were generated with anatomically constrained tractography (maximum length: 200 mm, cut off for FOD amplitude: 0.06), setting the boundary between the white and gray matter as seeds. These streamlines were refined from 100,000,000 streamlines to 10,000,000 streamlines with Spherical-deconvolution Informed Filtering of Tractograms (SIFT) to alleviate the overestimation of the number of streamlines. Commissural fibers connecting between each identical bilateral area defined by M132 Lyon macaque brain atlas (Markov et al., 2014) were extracted. Finally, automated fiber-tract quantification (https://github.com/yeatmanlab/AFQ) (Yeatman et al., 2012) was used to exclude outliers within each bundle (i.e., tracts that deviate more than three standard deviations (SD) from the mean location of each bundle), and to allocate 20 equidistant points on each tract. The bundles with more than five streamlines for each macaque were utilized for further analysis. ΔR_2_* and the other microstructural property on each point was calculated to obtain the median value for each point of each bundle and was averaged across macaques.

To investigate the relationship between vascularity and white matter microstructure, we estimated microstructural properties from dMRI using the Neurite Orientation Dispersion and Density Imaging (NODDI) toolbox (version 1.0.3; www.nitrc.org/projects/noddi_toolbox) (Zhang et al., 2012). NODDI models three compartments (intra-neurite, extra-neurite, and cerebrospinal fluid), using multi-shell high-angular resolution dMRI data (Zhang et al., 2012). Default parameter values were used for model fitting, including an intrinsic free diffusivity of 1.7 μm^2^/ms and an isotropic diffusivity of 3.0 μm^2^/ms. Two indices were used for further analysis: the neurite density index (NDI), representing the density of neurite compartments, and the orientation dispersion index (ODI), reflecting the dispersion of neurite orientations. FSL’s *dtifit* was applied to estimate the principal diffusion direction in each voxel. This was done solely for visualization purposes rather than for detailed modeling of fiber architecture.

### 2.4 Statistical Analysis

The group-average ΔR_2_* values between the gray and superficial white matter were compared using a two-tailed paired two-sample t-test. ΔR_2_* values from EL4a were used to represent the gray matter. To assess how inter-areal variability in ΔR_2_* of the superficial white matter is influenced by the overlying gray matter, we calculated the Pearson’s correlation coefficient between ΔR_2_* values of the gray matter and superficial white matter. To corroborate the relations between ΔR_2_* and microstructural properties within each area, Pearson’s correlation coefficients were calculated between ΔR_2_* and T1w/T2w-FLAIR (myelin map). We also calculated the coefficients with neuron density and receptor density in the overlying gray matter of each area to evaluate how neuron and synapse affect the vasculature in the superficial white matter. The myelin map corrected for B_0_ orientation bias was taken from our previous study (Autio et al., 2024), while neuron and receptor densities were from the literature (https://balsa.wustl.edu/study/P2Nql; (Froudist-Walsh et al., 2023)). These maps provide cortical area-wise, layer-averaged values derived from ex vivo histology and autoradiography. Consistent with our previous study (Autio et al., 2025), we focused on the total receptor density rather than individual receptor densities, as these were highly correlated with each other. To test whether these Pearson’s correlation coefficients differed between the gray matter and superficial white matter, we applied two-tailed bootstrap testing to each of these coefficients.

To examine vessel orientation characteristics in the superficial and deep white matter, we investigated the relationship between B_0_ orientation bias and the alignment of the cortex using B_0_-orientation uncorrected ΔR_2_* data. The vessel B orientation bias is defined by the following equation (Ogawa et al., 1993):

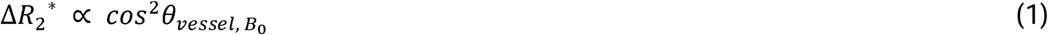

where, θ*_vessel, B0_* represents the angle between the primary orientation of the vessels within a voxel and the B_0_ direction in the individual’s original MRI space. In the superficial white matter, vessel alignment relative to the normal of cortex was indirectly assessed by calculating Pearson’s correlation coefficients between ΔR_2_^*^ and *cos*^2^θ*normal of cortex, B*_0_.

To model changes in ΔR_2_^*^, NDI, and ODI from the superficial to deep white matter, we fitted the group-average values of each metric against the relative distance from the superficial white matter to the midline (i.e., 0% distance at the white matter surface and 100% at the corpus callosum midline). Modeling was performed using a parsimonious exponential function:

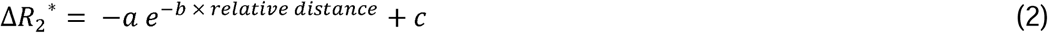

using the *fitnlm* function in Matlab R2016b (MathWorks). This empirical approximation was chosen to reflect gradual depth-dependent changes in the measured metrics, based on physiological considerations such as the exponential distance rule observed in axonal connectivity (Ercsey-Ravasz et al., 2013), quadratic relation between axon radius and energy consumption (Perge et al., 2012), cumulative axonal transport burden (Yang et al., 2023), and hierarchical arterial branching (Murray, 1926; Razavi et al., 2018). On the other hand, superficial white matter contains a mixture of smaller, more numerous axons (e.g., U-fibers) with more complex geometries (fanning, bending, and crossing), which inflates the ODI and decreases (apparent) NDI, and lower in deep white matter such as corpus callosum, where long-range fibers are coherently aligned. To evaluate this choice more rigorously, we compared the exponential model against the alternative models (linear, quadratic, and logistic) using corrected Akaike Information Criterion (AICc) and Bayesian Information Criterion (BIC). In addition, we evaluated the relationship between ΔR_2_^*^ and NDI, ODI, or path length across the white matter bundles using the Pearson’s correlation coefficient.

To determine whether ΔR_2_^*^ at each location could be predicted by fiber density, tract dispersion, and distance from superficial white matter to the midline (e.g., arteries branching or weakening toward deeper white matter), we used a forward stepwise nonlinear regression model. Here, we assumed a nonlinear relationship between the relative distance from the superficial white matter to the midline and ΔR_2_^*^, while treating the other properties as linear. Model selection was guided by the Bayesian information criterion. We evaluated the prediction accuracy using a leave-one-dataset-out (i.e., one white matter bundle out) cross-validation method. The significance of each Pearson’s correlation coefficient was assessed using two-tailed bootstrap testing. All statistical analyses were conducted using the Statistics and Machine Learning Toolbox in Matlab R2016b (MathWorks), with a significance threshold of *P* < 0.05. All P-values were Bonferroni-corrected for multiple comparisons.

## 3. Results

### 3.1 Charting Superficial White Matter Vascularity using Ferumoxytol-weighted MRI

The gradient map of mean ferumoxytol-weighted gradient-echo images provided clear visualization of macrovessels in superficial white matter (Fig. 1). Specifically, the ferumoxytol-weighted images enabled robust identification of macrovessels projecting from the pial surface and traversing along the superficial white matter (Fig. 1). These vessels likely correspond to those categorized as Durney’s Group 5 or 6 arteries as these vessels did not exhibit noticeable contrast in the pre-ferumoxytol images (Duvernoy et al., 1981).

**Figure 1.**
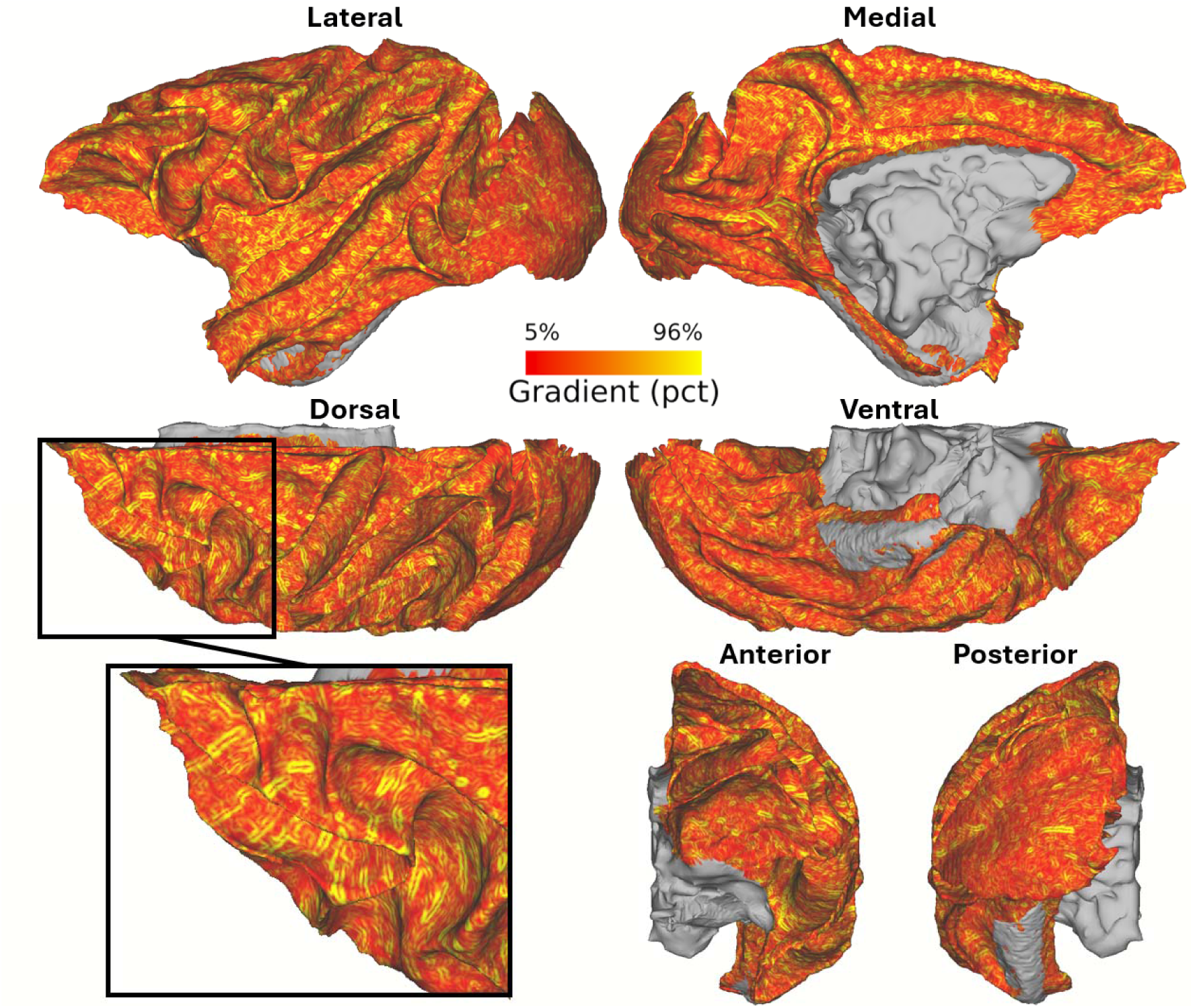
Charting macrovessels in the superficial white matter using ferumoxytol-weighted vessel-density informed MRI. Circular gradients represent vessels oriented perpendicular to the superficial white matter, while rod-like gradients denote vessels traversing along the superficial white matter. Gradients were notably stronger in the dorsal regions of the superficial white matter compared to the ventral regions. The black box indicates the location of the magnified view.

Comparison of vessel orientation in cortical gray matter and superficial white matter is illustrated in Figure 2 using spatial gradients of the mean ferumoxytol-weighted image. In the gray matter, the mean-intensity gradients exhibited ring-shaped patterns (Fig. 2A), indicative of vessels traversing perpendicular to the surface of the cerebral cortex (Autio et al., 2025). Conversely, in the superficial white matter, the gradients exhibited rod-shaped patterns (Fig. 2C) indicative of vessels traversing parallel to the white matter surface. These distinct vessel orientations are in good agreement with brain vessel anatomy (Duvernoy et al., 1981), and support Viessmann and colleagues’ indirect MRI observations of vessel orientation differences between gray and white matter (Viessmann et al., 2022). The sharp contrast was observed in orientation of vessels between equidistant superficial white matter (Fig. 2C) and the innermost EL of cortical gray matter (EL6b; Fig. 2B), supporting that the superficial white matter boundary was precisely delineated.

**Figure 2.**
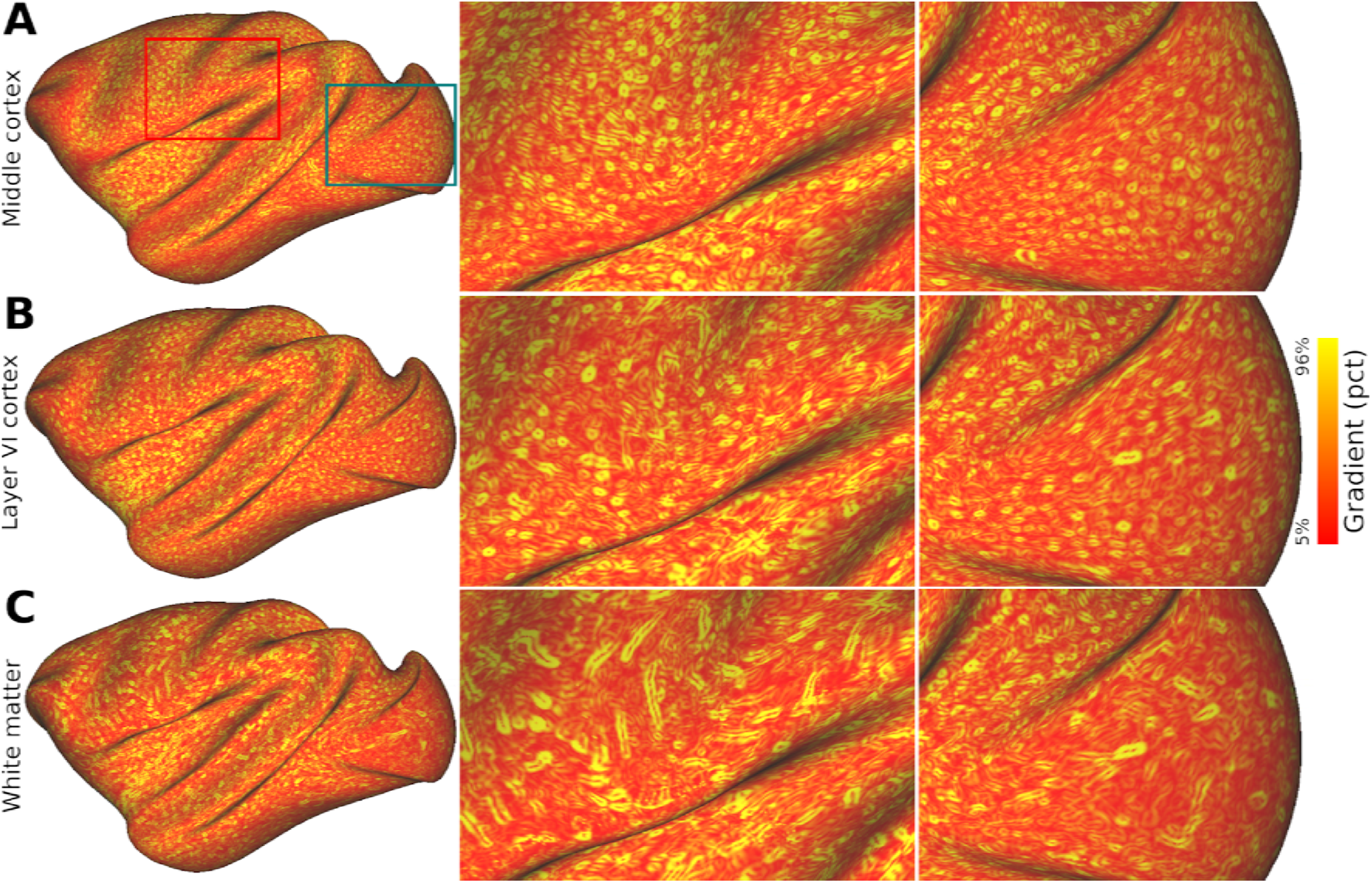
Distinct macrovessel arrangement in gray and white matter. Spatial gradients of the ferumoxytol-weighted image on the equivolumetric layer (EL) **(A)** EL4a, located proximal to cortical midthickness, and **(B)** EL6b, located adjacent to the white matter surface. **(C)** Equidistant superficial white matter surface located beneath the cortical gray matter. Note the striking contrast in the gradient pattern between the gray matter (circular structures) and superficial white matter (rod-like structures): circular gradients indicate the presence of vessels oriented perpendicular to the orientation of the cerebral cortex whereas rod-like gradients indicate vessels traversing along the white and gray matter interface. The red and green boxes indicate the locations of zoomed views.

We also observed areal differences in vascularization patterns. For instance, in V1 we observed sparse distribution of macrovessels in superficial white matter, contrasting the more uniform and dense distribution in the gray matter. In the prefrontal cortex, superficial white matter vessels exhibited orthogonal orientations whereas less distinct and more subtler variations in vessel orientation were observed near the temporal pole (Supp. Fig. 1).

Quantitative analysis revealed significant differences in vessel orientation bias between gray matter and superficial white matter. In the gray matter, correlations between ΔR_2_* and cerebral cortex orientation (cos^2^ θ in Eq. (1)) were negative (r = -0.32 – -0.26, t_3_ = -20.25, *P* = 0.0003), indicating that larger vessels predominantly align perpendicularly to the cortical surface. Conversely, the superficial white matter showed positive correlations (r = 0.05 – 0.11; t_3_ = 6.69, *P* = 0.0068), indicating parallel orientation of vessels relative to the cortical surface (Supp. Fig. 2).

We quantified vascularity in the superficial white matter by measuring R_2_* before and after ferumoxytol contrast agent injection, using ΔR_2_* as a well-established proxy for vascular volume (Kim et al., 2013). To mitigate the influence of macrovessels on ΔR_2_* (Fig. 1), we ascribed median ΔR_2_* to each M132-atlas parcel (Fig. 3A, upper panel) (Autio et al., 2025). While occipital and temporal gray matter exhibited a decrease in ΔR_2_* from lower-to higher-order areas (Autio et al., 2025; Felleman and Van Essen, 1991), no such trend was found in the superficial white matter. Interestingly, ΔR_2_* increased gradually from posterior to anterior regions within the frontal superficial white matter (Fig. 3A, bottom panel), a pattern absent in gray matter. Superficial white matter vascularization was not significantly correlated with that of gray matter (r = -0.10, *P* = 0.34; Fig. 3C), consistent with distinct energy demands between these regions (Harris and Attwell, 2012). We found a small, but significant, correlation with cortical curvature (r = 0.07, *P* < 0.001) and sulcal depths (r = 0.22, *P* < 0.001), revealing that there is a small geometrical effect influencing superficial white matter vasculature (Supp. Fig. 3). Although ΔR_2_* was systematically lower in superficial white matter compared to gray matter (Fig. 3B; paired one-sample t-test, t_87_ = 22.2, *P* < 0.001), variability in superficial white matter vascularization was comparable to gray matter (gray matter: SD = 3.56, interquartile range [IQR] = 4.00; superficial white matter: SD = 3.72, IQR = 4.68; F_87,87_ = 0.92, *P* = 0.69).

**Figure 3.**
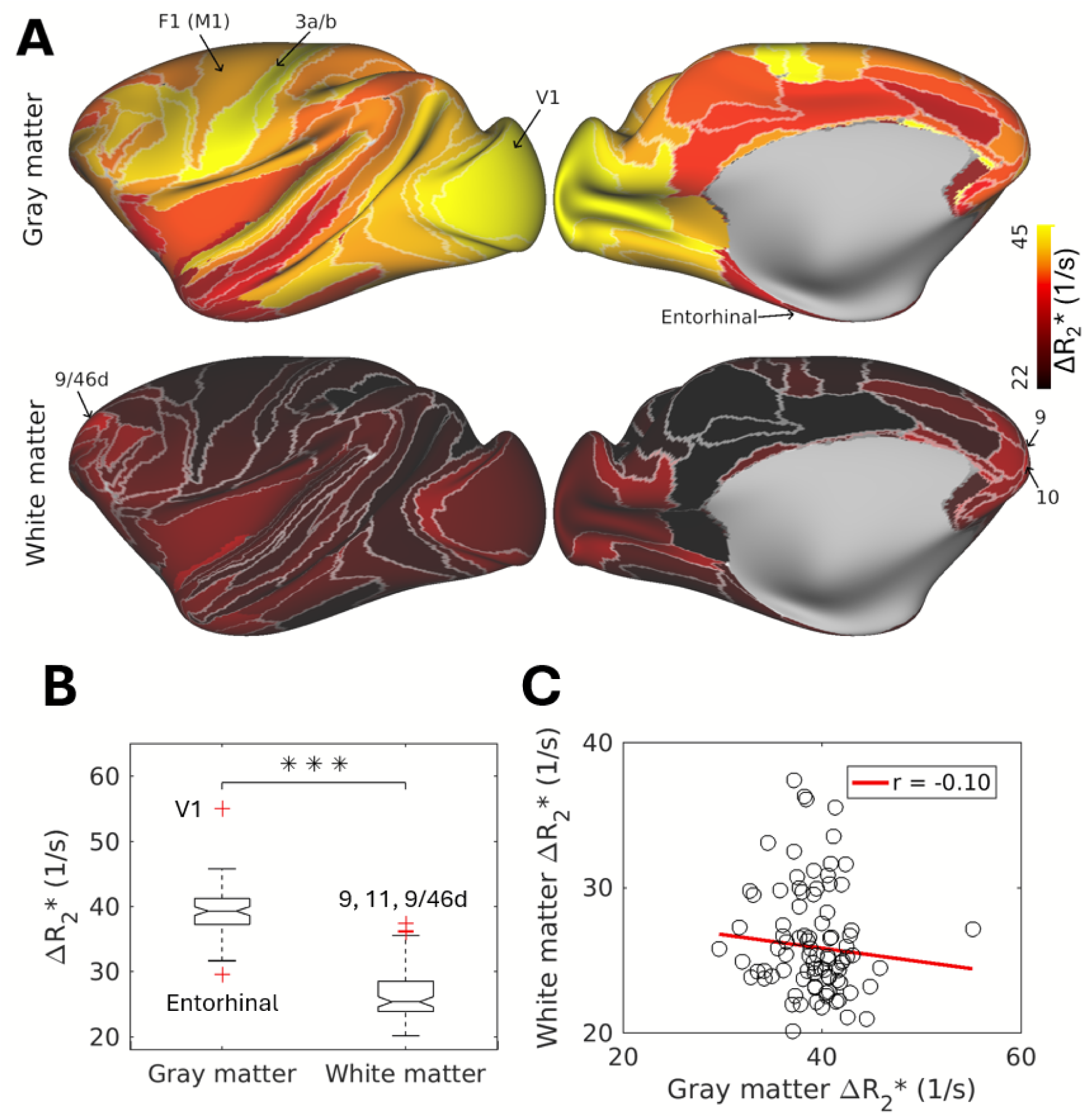
Comparison of vasculature in cortical gray matter and superficial white matter. **(A)** Vascular volume in cortical gray matter (upper panel) and superficial white matter (lower panel), estimated using ferumoxytol-induced change in transverse relaxation rate (ΔR_2_*). To reduce the influence of macrovessels, data was parcellated using the M132 areal atlas (Markov et al., 2014), and median ΔR_2_* value was assigned to each cortical area. **(B)** Box plot showing comparison of ΔR_2_* values between cortical gray matter and superficial white matter. Red crosses indicate outliers: in gray matter V1 and entorhinal cortex, and in superficial white matter areas 9, 11 and 9/46d. *** *P* < 0.001. **(C)** Scatter plot between vascular volume in superficial white matter and cortical gray matter (r = -0.01, *P* = 0.34).

### 3.2 Comparison between Superficial White Matter Vasculature and Overlying Cortical Neuron-Synapse Architecture

Given the large inter-areal heterogeneity in superficial white matter vasculature, we sought to investigate how neuroanatomical factors in the white matter and overlying cortical gray matter might influence superficial white matter energy consumption. To address this, we examined correlations between white matter ΔR_2_* and microstructural metrics, including neuron, myelin, and synaptic densities (Fig. 4) (Autio et al., 2025; Collins et al., 2010; Froudist-Walsh et al., 2023). In the cortical gray matter, ΔR_2_* demonstrated significant positive correlations with neuron density (r = 0.53, *P* < 0.001) and myelin density (r = 0.54, *P* < 0.001) and a negative correlation with receptor density (r = -0.42, *P* < 0.001) (Autio et al., 2025) (Fig. 4A–D, G). Conversely, within the superficial white matter, ΔR_2_* was positively correlated with receptor density (r = 0.50, *P* < 0.001) and negatively correlated with myelin density (r = -0.31, *P* < 0.001) (Fig. 4A, C, E–G), with no significant correlation observed for neuron density (r = -0.10, *P* = 0.36). Bootstrap testing confirmed significant differences between these correlations in the cortical gray matter and superficial white matter across all metrics (all *P* < 0.001). The M132-atlas parcellated ΔR_2_* also showed a significant positive correlation with myelin (r = 0.67, *P* < 0.001) within the cortical gray matter and a significant negative correlation with myelin (r = -0.28, *P* = 0.0014) within the superficial white matter (Supp. Fig. 4).

**Figure 4.**
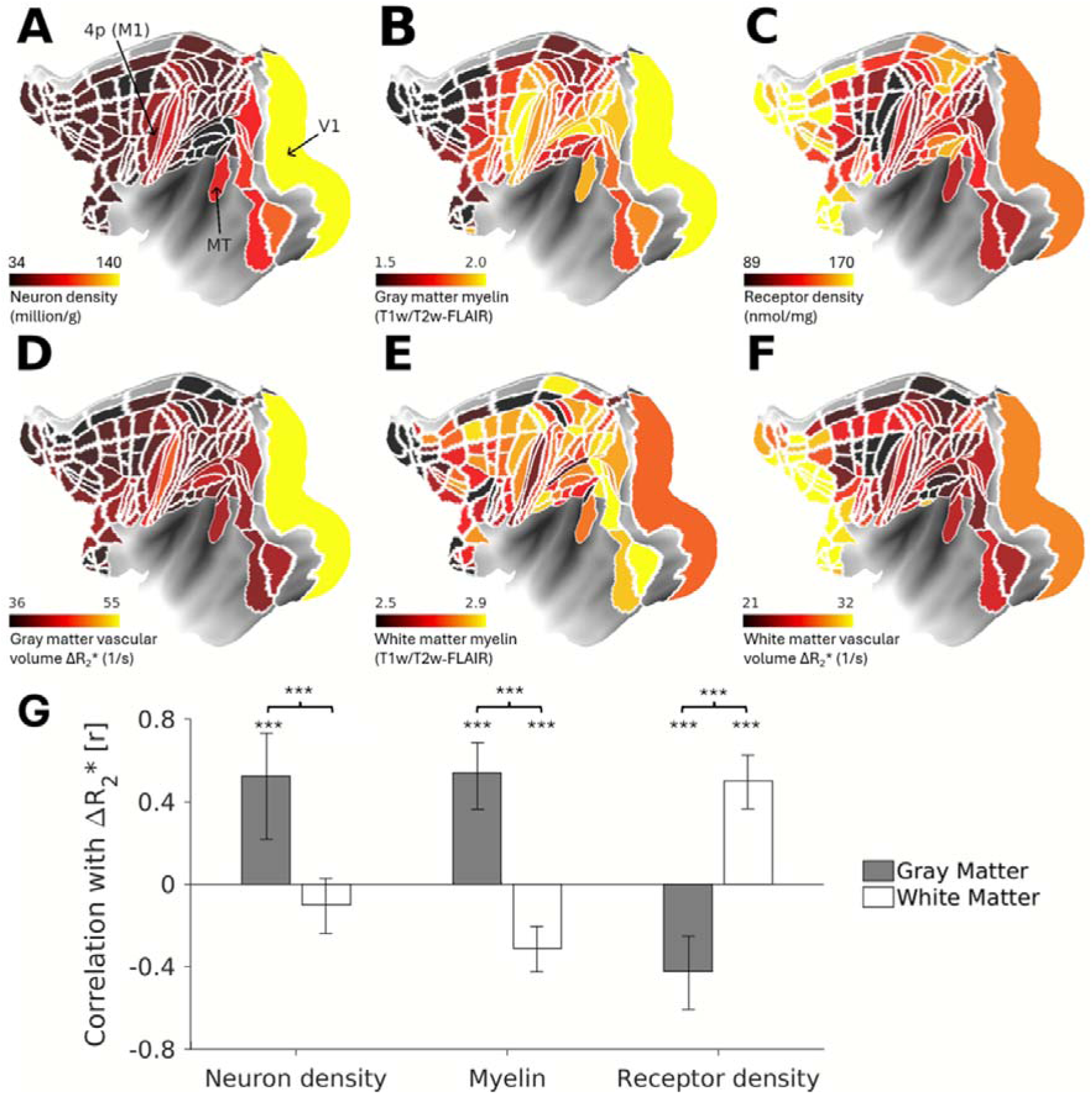
Neuroanatomical correlates of superficial white matter vascularity. Parcellated maps of cortical **(A)** neuron density (Collins et al., 2010), **(B)** T1w/T2w-FLAIR myelin index (Autio et al., 2024), **(C)** receptor density (Froudist-Walsh et al., 2023), **(D)** vascular volume assessed using ferumoxytol-induced change in transverse relaxation rate (ΔR_2_*) (Autio et al., 2025). Superficial white matter **(E)** T1w/T2w-FLAIR myelin index and **(F)** vascular volume. **(G)** Correlation between vascular volume and neuroanatomical factors. *** *P* < 0.001 (Bonferroni-corrected).

### 3.3 Comparison between White Matter Density, Orientation and Vasculature

Finally, we explored the vascularity deep in the white matter using vessel-density informed ferumoxytol-weighted MRI. To delineate the vessels in the volume-space, we applied a Hessian-based Frangi ‘vesselness’ filter to the TE-averaged mean intensity images (Frangi et al., 1998). This analysis revealed a large number of vessels traversing from the pia mater to white matter (2800 – 4220 mm^3^, 13–15 % of white matter voxels; Fig. 5A). Visual inspection of these vessels revealed that several vessels projected toward lateral ventricles, likely corresponding to the large draining veins. Several vessels appeared to traverse along the major tracts (e.g., commissural fibers connecting brain hemispheres), raising the question whether the feeding arterial vessels are organized along the major tract bundles (Smirnov et al., 2021).

**Figure 5.**
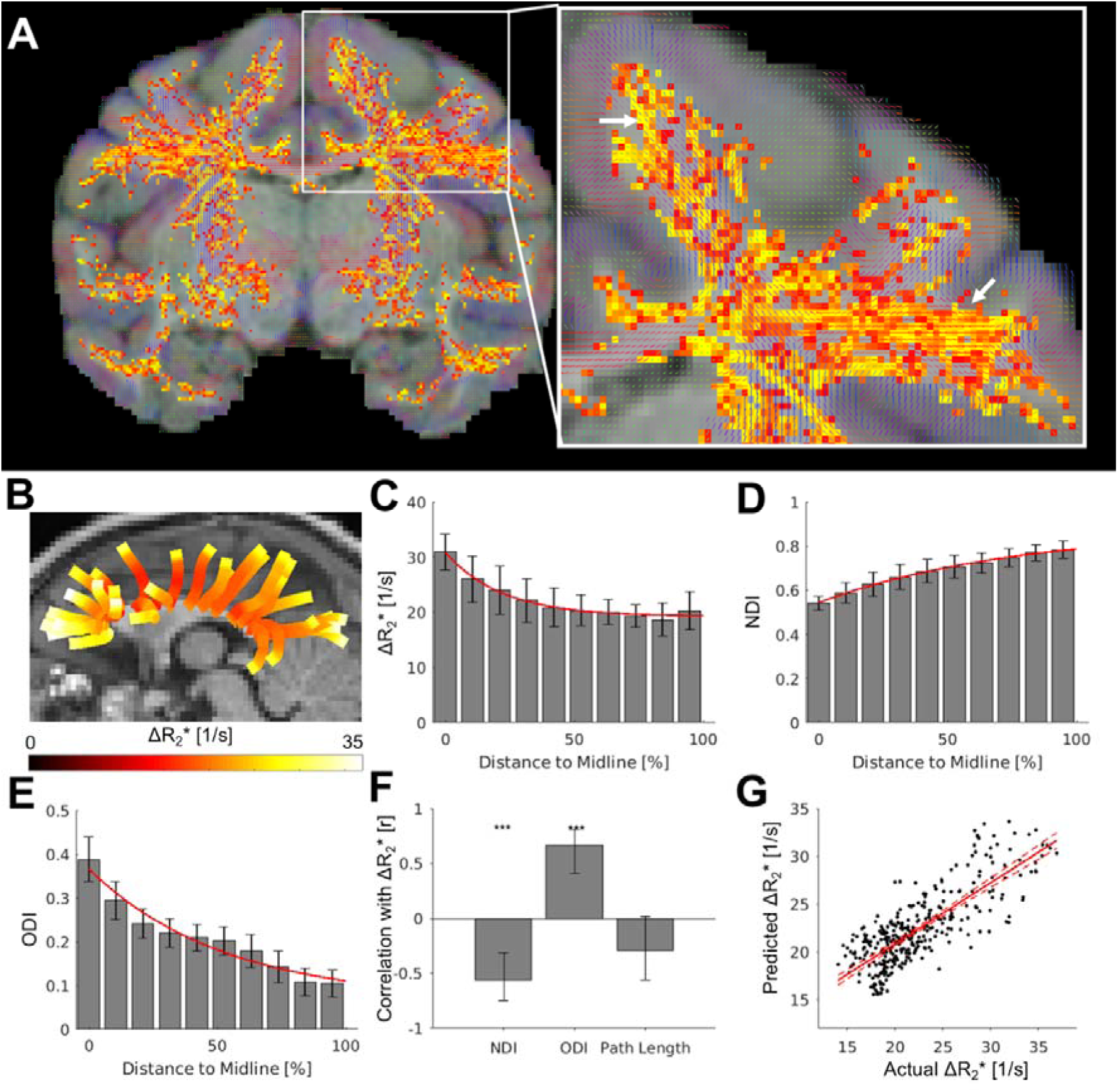
Modeling white matter vascularity using tract density, dispersion and distance from cortical surface. **(A)** Diffusion tensor orientation (red, blue, and green color indicate the orientation of major fibers are mainly in right-left, superior-inferior, and anterior-posterior, respectively) overlaid with maximum intensity projection image of ferumoxytol-weighted and frangi-filtered vessel maps (red-yellow). **(B)** Diffusion tractography. **(C)** ferumoxytol-induced change in transverse relaxation rate (ΔR_2_*), **(D)** neurite density index (NDI), and **(E)** orientation dispersion index (ODI) values calculated from white matter surface (0%) toward the cortical midline (100%). **(F)** Pearson’s correlation coefficients between ΔR_2_* and NDI, ODI, or path length across tracts. **(G)** Relationship between the ΔR_2_* within the white matter and the value predicted by the microstructural factors. Solid red line indicates the fit by the linear model, while dashed lines indicate 95% confidence interval. *** *P* < 0.001 (Bonferroni-corrected).

To corroborate this question, we compared vessel orientation using a Frangi-filter and assessed fiber tract orientation estimated by diffusion tensor analysis (Basser et al., 1994). Our visual inspection revealed that, in most cases, the vessels were aligned with the primary orientation of axons within the commissural fiber bundles (Fig. 5A; white arrows). However, the spatial resolution of dMRI (0.9 mm isotropic; 0.73 mm^3^) was 60-fold lower than that of vessel imaging (0.23 mm isotropic; 0.012 mm^3^), making it challenging to accurately associate individual white matter bundles with their corresponding vessels, particularly smaller ones.

To overcome the spatial resolution limitation, we compared vascular density with NODDI, a model that estimates neurite density and orientation dispersion within each white matter voxel (Zhang et al., 2012). Specifically, we focused on commissural fibers connecting identical bilateral regions, as these long bundles can be robustly delineated (Fig. 5B). Within each bundle, as the relative distance from the white surface to the midline increased, ΔR_2_* and ODI exhibited exponential decay (ΔR_2_*: _11.64 e_-0.05 × relative distance _+19.13; ODI: 0.30 e_-0.02 × relative distance _+ 0.07), whereas NDI_ showed exponential growth (−0.32 e^-0.01^ ^×^ ^relative^ ^distance^ + 0.87) (Fig. 5C-E; for the relationship with raw axon path length, see Supp. Fig. 5). The data supported the exponential model over linear, quadratic and logistic models (Supp. Table 1). Across all bundles, ΔR_2_* was significantly negatively correlated with NDI (r = -0.51, *P* < 0.001) and positively correlated with ODI (r = 0.67, *P* < 0.001), while it was not significantly correlated with path length (r = -0.27, *P* = 0.32).

Given the systematic variation between microstructure, brain geometry and vasculature, we additionally asked whether the white matter microstructure may predict the variation in white matter vasculature. To address this question, we performed a forward stepwise nonlinear regression model selected by Bayesian information criterion (see methods). This analysis revealed that ΔR_2_* along each bundle can be predicted by the NDI, ODI, and the relative distance from the white matter surface to the midline (corpus callosum) with high precision (r = 0.79, *P* < 0.001; Fig. 5G).

## 4. Discussion

In this study, we explored the relationship between neuron-axon-synapse architecture and the organization of white matter vasculature using high-resolution multimodal MRI and quantitative anatomical data in a non-human primate brain. Our novel surface presentation of superficial white matter vasculature revealed that the density of cortical synapses and axons, rather than neurons, significantly influences the white matter vascularity. In deep white matter, we found that the geometric arrangement of white matter bundles, neurite density, and the distance from the cortical surface are key determinants of white matter vascular organization. These findings represent both a methodological and conceptual advancement in white matter vasculature MRI, offering potential clinical applications in understanding aging-related white matter pathologies.

### 4.1 Heterogeneity of Superficial White Matter Vasculature

The superficial white matter macrovasculature is a largely unexplored area with limited studies (Duvernoy et al., 1981; Hase et al., 2019; Smirnov et al., 2021). It displays a complex and heterogeneous organization with several notable organizational features. In regions where white matter blades are exceptionally thin (< 0.5 mm), such as underneath V1, spatial constraints confine vessel trajectories to the white matter surface. Certain vessels take the most direct route to the lateral ventricles, possibly reflecting preferential venous drainage patterns (Smirnov et al., 2021). Additionally, some of the macrovessels exhibit orthogonal orientations, appearing to cross each other (Supp. Fig. 1; arrows). Additionally, some of the macrovessels exhibit orthogonal orientations, appearing to cross each other (Supp. Fig. 1; arrows). These vessels may be the arteries that deliver oxygen and nutrients to cortical layer VI (Duvernoy et al., 1981). Hypoperfusion or ischemia in the superficial white matter vasculature may selectively disrupt energy supply to cortical layer VI, potentially impairing corticothalamic and corticoclaustral circuits (Wiser and Callaway, 1996).

One surprising discovery of the study was the substantial two-fold variability in the superficial white matter vasculature (Fig. 3A, B). This heterogeneity likely reflects a number of factors, including the volume of traversing axonal tracts, hierarchical patterns of arterial branching (Razavi et al., 2018), and the diverse functional properties of different white matter regions. For instance, axonal conduction velocities vary three-fold (3.1 – 9.4 m/s) (Innocenti et al., 2014), and the frequency bands involved in inter-areal communication range from theta to gamma band (Shu et al., 2003; Vezoli et al., 2021). Such functional diversity is enabled by differences in axon diameter and myelination, influencing local energy requirements and vascular densities (Perge et al., 2012). However, we found negligible correlation between the vascular volume in gray matter and superficial white matter (Fig. 3C), implying that the metabolic demands are driven by distinct biophysical mechanisms. This provides experimental validation to the previous modeling study (Harris and Attwell, 2012). That study suggested that, while energy demand in gray matter is primarily influenced by the frequency of electrical activity (e.g., Na^+^-K^+^ ATPase of neuronal cell membrane potential), in white matter it is primarily determined by distinct biophysical processes related to the maintenance of axons and their surroundings.

Another unexpected finding was that an indirect proxy measure of myelin density, as assessed using T1w/T2w-FLAIR, accounted for only about 10% of the variance in the superficial white matter vasculature (Fig. 4G). This result is surprising given that myelin is generally considered a key factor in reducing the energetic demands of axonal conduction (Saab et al., 2013), yet it aligns with biophysical simulation by (Harris and Attwell, 2012), which suggest that the net energy savings from myelination are modest when factoring in the total energy demands of myelin synthesis and maintenance.

Interestingly, myelin was positively correlated with the gray matter vasculature and negatively with the white matter vasculature. This dissociation likely reflects fundamental functional differences between the two tissue types. In the gray matter, myelin reduces the refractory period (Nashmi and Fehlings, 2001), enabling higher-frequency action potentials that elevate metabolic demand. Additionally, myelin inhibits synaptogenesis (Froudist-Walsh et al., 2023), and neuronal somata, which are more dense in myelin-rich regions, consume more energy per unit volume than the surrounding neuropil (Autio et al., 2025; Wong-Riley, 1989). In contrast, in white matter, myelin reduces energy usage by insulating axons and suppressing collateral branching into mitochondria-rich segments (Filbin, 2003; Tao et al., 2014). These contrasting roles support the view that the dominant sources of metabolic demand differ between the gray and white matter (Harris and Attwell, 2012). However, we acknowledge that the utility of T1w/T2w-FLAIR as a biomarker of myelin may be more limited in the white matter than cortical gray matter (Glasser et al., 2014; Sandrone et al., 2023). Thus, further histological investigations are essential to corroborate how myelin, and its supporting cells, such as oligodendrocytes, influence white matter vascularization (Biswas et al., 2020; Gao et al., 2023).

### 4.2 Cortical Synaptic Architecture Shapes Superficial White Matter Vasculature

Given that axons originate from neuronal cell bodies, and growth factors guide the alignment of axons and arteries outside the brain (Carmeliet and Tessier-Lavigne, 2005), we initially hypothesized that white matter vascularization would correlate with cortical neuron density. However, we found that the vascular density in the superficial white matter was not significantly associated with neuron density (i.e., axon origin) (Fig. 4G). Instead, it showed a positive correlation with receptor density (i.e., axon terminals) in the overlying cortical gray matter (Froudist-Walsh et al., 2023). This suggests that axonal branches may drive local metabolic demand, possibly due to their high mitochondrial density (Tao et al., 2014), rather than by the total number of axons stemming from cortical soma.

Receptor density, however, does not distinguish between pre- and post-synaptic components and therefore cannot differentiate whether receptors are located on axon terminals or dendrites. In cortical areas, the majority of receptors are localized on dendrites, particularly of pyramidal neurons. As highlighted by (Froudist-Walsh et al., 2023), receptor density strongly covaries with dendritic complexity, and pyramidal dendritic arborization increases along a gradient from sensorimotor to association areas. These extensive dendritic trees are critical for integrating synaptic inputs from both local and long-range afferents. Importantly, increased dendritic complexity in association areas often co-occurs with larger projection neurons, which possess greater soma sizes (Cullheim, 1978; Lee et al., 1986), enabling the biosynthetic support required for maintaining long, thick, and myelinated axons.

Thus, the observed correlation between receptor density and superficial white matter vascularization may reflect an indirect relationship (Fig. 4C, F, G): areas with more complex dendritic trees and higher receptor densities tend to contain projection neurons that not only integrate more inputs but also send long-range axonal outputs, increasing metabolic demand in both gray and white matter compartments. This dual demand — on dendrites for input integration and on axons for long-distance transmission — may drive the need for enhanced vascular support in the underlying white matter. Nonetheless, there are exceptions to this general pattern. For example, regions such as the cingulate cortex exhibit extensive dendritic arbors yet relatively small somata and relatively short inter-areal connections (Elston et al., 2005; Markov et al., 2013), suggesting that receptor density alone cannot fully account for white matter vascular variation.

Additionally, axonal transport may contribute to the relationship between cortical receptor density and superficial white matter vascularization (Fig. 4G), as well as to the observed vascular gradient from superficial white matter toward the corpus callosum. Both anterograde (from cell body toward synapse) and retrograde (from synapse toward cell body) transport are essential for delivering and recycling mitochondria, organelles, proteins, and signaling molecules along the axon, including branch points, internodes, and nodes of Ranvier. These active transport processes are energetically demanding, particularly through maintenance of the actin cytoskeleton (Fath and Lasek, 1988; Harris and Attwell, 2012), and may contribute significantly to white matter energy consumption. Because all metabolic substrates originate from the cortex, the energetic burden of transport is cumulative: the longer the axon, the greater the metabolic cost. This gradient in transport demand may contribute to increased vascular support near the cortex and could contribute to the exponential decay of axonal connection length with distance, as described by the distance rule (Ercsey-Ravasz et al., 2013). Considering that receptors are directly associated with retrograde transport, the greater number of axon terminals may facilitate large-scale retrograde transport, thus increasing energy consumption and vessel density. However, the proportion of energy devoted to anterograde and retrograde transport remains debated, with estimates ranging from 2% to 50% of total white matter energy consumption (Bernstein and Bamburg, 2003; Daniel et al., 1986; Engl and Attwell, 2015; Guedes-Dias and Holzbaur, 2019; Harris and Attwell, 2012; Yang et al., 2023), and more advanced research designs are needed to resolve the energy dynamics of these processes.

Another potential factor is white matter plasticity (Kimura et al., 2024; Sampaio-Baptista and Johansen-Berg, 2017). Increased neural synchronization (between cortical regions) may induce microstructural changes within white matter (Lazari et al., 2022; Sampaio-Baptista et al., 2021), such as myelination or oligodendrogenesis (Gibson et al., 2014). Given that synchronization also increases receptor densities through spike-timing-dependent plasticity (Caporale and Dan, 2008; Lamsa et al., 2010), high receptor density may indicate frequent plasticity in the underlying white matter, leading to increased energy demand.

### 4.3 White Matter Vasculature Reflects Fiber Architecture

Our study also revealed that macrovessels tend to align with the major orientation of axons. This supports previous human MRI studies that showed B_0_ orientation bias in ΔR_2_* (Hernández-Torres et al., 2017) and T_2_*-weighted signal-intensity (Viessmann et al., 2022). The observed alignment may reflect common developmental mechanisms, such as shared guidance cues involving neuropilin-plexin signaling, which regulate both axons and arterial vascular patterning (Carmeliet and Tessier-Lavigne, 2005). However, very recent susceptibility-weighted imaging (SWI) findings show that venous vessel alignment with axons is not uniform across brain regions (Schilling et al., 2025).

The alignment of feeding arteries with fiber tracts may in part explain why blood volume is higher in regions where multiple white matter tracts intersect, such as beneath multimodal association areas that form a dense interconnected network (Fig. 5B) (Markov et al., 2014). This axonal-vascular relationship is influenced by anatomical constraints and may also relate to variations in axon diameter (Fig. 5F). Axons with larger diameters transmit signals faster and require more energy to maintain their function, which could result in increased vessel density to support the higher energy demands (Harris and Attwell, 2012; Perge et al., 2012). In contrast, smaller diameter axons, which constitute the majority of axons in the central nervous system, conduct signals more slowly but are more energy-efficient (Perge et al., 2012). Given that high NDI may reflect small axon diameter in that region (Alexander et al., 2010), the negative relationships between vessel density and NDI may be explained by the energy demand defined by axon diameter.

Overall, these findings highlight the complex interplay between anatomical structures that influence the vascularization of white matter. Indeed, a forward stepwise nonlinear regression model selected by Bayesian information criterion predicts white matter vascularity with high precision (R^2^ = 0.62) (Fig. 5G). This information could be valuable for modeling brain blood circulation, understanding white matter pathology, and guiding neurosurgical operations. Moving forward, it will be fundamental to augment vascular models with arterial and venous classifications —more feasible at ultra-high field strengths (Caan et al., 2019) — as well as more advanced dMRI modeling approaches (Coelho et al., 2022; Jbabdi et al., 2012; Tahedl et al., 2025; Tournier et al., 2019) and quantitative anatomical data acquired in the same individuals.

## 5. Conclusions

This study examined the relationship between neuron-axon-synapse architecture and white matter vasculature using high-resolution multimodal MRI and quantitative anatomy in primates. We found that white matter vasculature was correlated to the synaptic density of the overlying gray matter, while myelinated tracts were characterized by sparser vascular networks. By integrating axon geometry, density, and distance from the cortical surface, a predictive model was developed to map vascularity, providing new insights into the structural and metabolic determinants of white matter vasculature.

## Data and Code Availability

The data for the figures will be made available at https://balsa.wustl.edu/study/show/LPDP.

These datasets require Connectome Workbench v2.0 for visualization.

The NHP-HCP pipelines are available at https://github.com/Washington-University/NHPPipelines.

## CRediT authorship contribution statement

**Ikko Kimura:** Conceptualization, Software, Formal analysis, Writing - original draft, Writing - review & editing, Visualization. **Takuya Hayashi:** Resources, Funding acquisition. **Joonas Autio:** Conceptualization, Methodology, Formal analysis, Writing - original draft, Writing - review & editing, Visualization, Resources, Funding acquisition.

## Acknowledgements

The authors appreciate discussions and technical contributions from Akiko Uematsu, Naofumi Yoshida, Takayuki Ose, Masahiro Ohno, and Reiko Kobayashi.

## References

1. Alexander, D.C., Hubbard, P.L., Hall, M.G., Moore, E.A., Ptito, M., Parker, G.J.M., Dyrby, T.B., 2010. Orientationally invariant indices of axon diameter and density from diffusion MRI. Neuroimage 52, 1374–1389. 10.1016/j.neuroimage.2010.05.043

2. Autio, J.A., Glasser, M.F., Ose, T., Donahue, C.J., Bastiani, M., Ohno, M., Kawabata, Y., Urushibata, Y., Murata, K., Nishigori, K., Yamaguchi, M., Hori, Y., Yoshida, A., Go, Y., Coalson, T.S., Jbabdi, S., Sotiropoulos, S.N., Kennedy, H., Smith, S., Van Essen, D.C., Hayashi, T., 2020. Towards HCP-Style macaque connectomes: 24-Channel 3T multi-array coil, MRI sequences and preprocessing. Neuroimage 215, 116800. 10.1016/j.neuroimage.2020.116800

3. Autio, J.A., Kimura, I., Ose, T., Matsumoto, Y., Ohno, M., Urushibata, Y., Ikeda, T., Glasser, M.F., Van Essen, D.C., Hayashi, T., 2025. Mapping vascular network architecture in primate brain using ferumoxytol-weighted laminar MRI. 10.7554/elife.99940.3

4. Autio, J.A., Roberts, R.E., 2014. Interpreting functional diffusion tensor imaging. Front. Neurosci. 8, 68. 10.3389/fnins.2014.00068

5. Autio, J.A., Uematsu, A., Ikeda, T., Ose, T., Hou, Y., Magrou, L., Kimura, I., Ohno, M., Murata, K., Coalson, T., Kennedy, H., Glasser, M.F., Van Essen, D.C., Hayashi, T., 2024. Charting cortical-layer specific area boundaries using Gibbs ringing attenuated T1w/T2w-FLAIR myelin MRI. bioRxiv. 10.1101/2024.09.27.615294

6. Basser, P.J., Mattiello, J., LeBihan, D., 1994. MR diffusion tensor spectroscopy and imaging. Biophys. J. 66, 259–267. 10.1016/S0006-3495(94)80775-1

7. Bernier, M., Chen, J.E., Ohringer, N., Fultz, N.E., Leaf, R.K., Viessmann, O., Lewis, L.D., Wald, L.L., Polimeni, J.R., 2019. Prospects for improving neuronal specificity of fMRI with ferumoxytol: an evaluation of vascular segmentation and cortical depth-dependent analysis, in: Proc Intl Soc Mag Reson Med. archive.ismrm.org, p. 537.

8. Bernstein, B.W., Bamburg, J.R., 2003. Actin-ATP hydrolysis is a major energy drain for neurons. J. Neurosci. 23, 1–6. 10.1523/JNEUROSCI.23-01-00002.2003

9. Biswas, S., Cottarelli, A., Agalliu, D., 2020. Neuronal and glial regulation of CNS angiogenesis and barriergenesis. Development 147. 10.1242/dev.182279

10. Bok, S.T., 1929. Der Einflu¥ der in den Furchen und Windungen auftretenden Krümmungen der Gro¥hirnrinde auf die Rindenarchitektur. Arch. Psychiatr. Nervenkr. Z. Gesamte Neurol. Psychiatr. 121, 682–750. 10.1007/BF02864437

11. Borowsky, I.W., Collins, R.C., 1989. Metabolic anatomy of brain: a comparison of regional capillary density, glucose metabolism, and enzyme activities. J. Comp. Neurol. 288, 401–413. 10.1002/cne.902880304

12. Boxerman, J.L., Hamberg, L.M., Rosen, B.R., Weisskoff, R.M., 1995. MR contrast due to intravascular magnetic susceptibility perturbations. Magn. Reson. Med. 34, 555–566. 10.1002/mrm.1910340412

13. Caan, M.W.A., Bazin, P.-L., Marques, J.P., de Hollander, G., Dumoulin, S.O., van der Zwaag, W., 2019. MP2RAGEME: T1 , T2 * , and QSM mapping in one sequence at 7 tesla. Hum. Brain Mapp. 40, 1786–1798. 10.1002/hbm.24490

14. Caporale, N., Dan, Y., 2008. Spike Timing–Dependent Plasticity: A Hebbian Learning Rule. 10.1146/annurev.neuro.31.060407.125639

15. Carmeliet, P., Tessier-Lavigne, M., 2005. Common mechanisms of nerve and blood vessel wiring. Nature 436, 193–200. 10.1038/nature03875

16. Chen, J.J., Rosas, H.D., Salat, D.H., 2013. The relationship between cortical blood flow and sub-cortical white-matter health across the adult age span. PLoS One 8, e56733. 10.1371/journal.pone.0056733

17. Chojdak-Łukasiewicz, J., Dziadkowiak, E., Zimny, A., Paradowski, B., 2021. Cerebral small vessel disease: A review. Adv. Clin. Exp. Med. 30, 349–356. 10.17219/acem/131216

18. Coelho, S., Baete, S.H., Lemberskiy, G., Ades-Aron, B., Barrol, G., Veraart, J., Novikov, D.S., Fieremans, E., 2022. Reproducibility of the Standard Model of diffusion in white matter on clinical MRI systems. Neuroimage 257, 119290. 10.1016/j.neuroimage.2022.119290

19. Collins, C.E., Airey, D.C., Young, N.A., Leitch, D.B., Kaas, J.H., 2010. Neuron densities vary across and within cortical areas in primates. Proceedings of the National Academy of Sciences 107, 15927–15932. 10.1073/pnas.1010356107

20. Cullheim, S., 1978. Relations between cell body size, axon diameter and axon conduction velocity of cat sciatic α-motoneurons stained with horseradish peroxidase. Neurosci. Lett. 8, 17–20. 10.1016/0304-3940(78)90090-3

21. Daniel, J.L., Molish, I.R., Robkin, L., Holmsen, H., 1986. Nucleotide exchange between cytosolic ATP and F-actin-bound ADP may be a major energy-utilizing process in unstimulated platelets. Eur. J. Biochem. 156, 677–684. 10.1111/j.1432-1033.1986.tb09631.x

22. Donahue, C.J., Sotiropoulos, S.N., Jbabdi, S., Hernandez-Fernandez, M., Behrens, T.E., Dyrby, T.B., Coalson, T., Kennedy, H., Knoblauch, K., Van Essen, D.C., Glasser, M.F., 2016. Using Diffusion Tractography to Predict Cortical Connection Strength and Distance: A Quantitative Comparison with Tracers in the Monkey. J. Neurosci. 36, 6758–6770. 10.1523/JNEUROSCI.0493-16.2016

23. Duvernoy, H.M., Delon, S., Vannson, J.L., 1981. Cortical blood vessels of the human brain. Brain Res. Bull. 7, 519–579. 10.1016/0361-9230(81)90007-1

24. Elston, G.N., Benavides-Piccione, R., Defelipe, J., 2005. A study of pyramidal cell structure in the cingulate cortex of the macaque monkey with comparative notes on inferotemporal and primary visual cortex. Cereb. Cortex 15, 64–73. 10.1093/cercor/bhh109

25. Engl, E., Attwell, D., 2015. Non-signalling energy use in the brain: Non-signalling energy use in the brain. J. Physiol. 593, 3417–3429. 10.1113/jphysiol.2014.282517

26. Ercsey-Ravasz, M., Markov, N.T., Lamy, C., Van Essen, D.C., Knoblauch, K., Toroczkai, Z., Kennedy, H., 2013. A predictive network model of cerebral cortical connectivity based on a distance rule. Neuron 80, 184–197. 10.1016/j.neuron.2013.07.036

27. Fath, K.R., Lasek, R.J., 1988. Two classes of actin microfilaments are associated with the inner cytoskeleton of axons. J. Cell Biol. 107, 613–621. 10.1083/jcb.107.2.613

28. Felleman, D.J., Van Essen, D.C., 1991. Distributed hierarchical processing in the primate cerebral cortex. Cereb. Cortex 1, 1–47. 10.1093/cercor/1.1.1-a

29. Filbin, M.T., 2003. Myelin-associated inhibitors of axonal regeneration in the adult mammalian CNS. Nat. Rev. Neurosci. 4, 703–713. 10.1038/nrn1195

30. Frangi, A.F., Niessen, W.J., Vincken, K.L., Viergever, M.A., 1998. Multiscale vessel enhancement filtering, in: Medical Image Computing and Computer-Assisted Intervention — MICCAI’98, Lecture Notes in Computer Science. Springer Berlin Heidelberg, Berlin, Heidelberg, pp. 130–137. 10.1007/bfb0056195

31. Froudist-Walsh, S., Xu, T., Niu, M., Rapan, L., Zhao, L., Margulies, D.S., Zilles, K., Wang, X.-J., Palomero-Gallagher, N., 2023. Gradients of neurotransmitter receptor expression in the macaque cortex. Nat. Neurosci. 26, 1281–1294. 10.1038/s41593-023-01351-2

32. Gao, L., Pan, X., Zhang, J.H., Xia, Y., 2023. Glial cells: an important switch for the vascular function of the central nervous system. Front. Cell. Neurosci. 17, 1166770. 10.3389/fncel.2023.1166770

33. Gibson, E.M., Purger, D., Mount, C.W., Goldstein, A.K., Lin, G.L., Wood, L.S., Inema, I., Miller, S.E., Bieri, G., Zuchero, J.B., Barres, B.A., Woo, P.J., Vogel, H., Monje, M., 2014. Neuronal activity promotes oligodendrogenesis and adaptive myelination in the mammalian brain. Science 344, 1252304. 10.1126/science.1252304

34. Glasser, M.F., Goyal, M.S., Preuss, T.M., Raichle, M.E., Van Essen, D.C., 2014. Trends and properties of human cerebral cortex: correlations with cortical myelin content. Neuroimage 93 Pt 2, 165–175. 10.1016/j.neuroimage.2013.03.060

35. Glasser, M.F., Sotiropoulos, S.N., Wilson, J.A., Coalson, T.S., Fischl, B., Andersson, J.L., Xu, J., Jbabdi, S., Webster, M., Polimeni, J.R., Van Essen, D.C., Jenkinson, M., WU-Minn HCP Consortium, 2013. The minimal preprocessing pipelines for the Human Connectome Project. Neuroimage 80, 105–124. 10.1016/j.neuroimage.2013.04.127

36. Goldstein, A.Y.N., Wang, X., Schwarz, T.L., 2008. Axonal transport and the delivery of pre-synaptic components. Curr. Opin. Neurobiol. 18, 495–503. 10.1016/j.conb.2008.10.003

37. Guedes-Dias, P., Holzbaur, E.L.F., 2019. Axonal transport: Driving synaptic function. Science 366, eaaw9997. 10.1126/science.aaw9997

38. Harris, J.J., Attwell, D., 2012. The energetics of CNS white matter. J. Neurosci. 32, 356–371. 10.1523/JNEUROSCI.3430-11.2012

39. Hase, Y., Ding, R., Harrison, G., Hawthorne, E., King, A., Gettings, S., Platten, C., Stevenson, W., Craggs, L.J.L., Kalaria, R.N., 2019. White matter capillaries in vascular and neurodegenerative dementias. Acta Neuropathol. Commun. 7, 16. 10.1186/s40478-019-0666-x

40. Hayashi, T., Hou, Y., Glasser, M.F., Autio, J.A., Knoblauch, K., Inoue-Murayama, M., Coalson, T., Yacoub, E., Smith, S., Kennedy, H., Van Essen, D.C., 2021. The nonhuman primate neuroimaging and neuroanatomy project. Neuroimage 229, 117726. 10.1016/j.neuroimage.2021.117726

41. Hernández-Torres, E., Kassner, N., Forkert, N.D., Wei, L., Wiggermann, V., Daemen, M., Machan, L., Traboulsee, A., Li, D., Rauscher, A., 2017. Anisotropic cerebral vascular architecture causes orientation dependency in cerebral blood flow and volume measured with dynamic susceptibility contrast magnetic resonance imaging. J. Cereb. Blood Flow Metab. 37, 1108–1119. 10.1177/0271678X16653134

42. Huang, Y., Wei, P.-H., Xu, L., Chen, D., Yang, Y., Song, W., Yi, Y., Jia, X., Wu, G., Fan, Q., Cui, Z., Zhao, G., 2023. Intracranial electrophysiological and structural basis of BOLD functional connectivity in human brain white matter. Nat. Commun. 14, 3414. 10.1038/s41467-023-39067-3

43. Huang, Y., Yang, Y., Hao, L., Hu, X., Wang, P., Ding, Z., Gao, J.-H., Gore, J.C., 2020. Detection of functional networks within white matter using independent component analysis. Neuroimage 222, 117278. 10.1016/j.neuroimage.2020.117278

44. Innocenti, G.M., Vercelli, A., Caminiti, R., 2014. The diameter of cortical axons depends both on the area of origin and target. Cereb. Cortex 24, 2178–2188. 10.1093/cercor/bht070

45. Jbabdi, S., Sotiropoulos, S.N., Savio, A.M., Graña, M., Behrens, T.E.J., 2012. Model-based analysis of multishell diffusion MR data for tractography: how to get over fitting problems. Magn. Reson. Med. 68, 1846–1855. 10.1002/mrm.24204

46. Kageyama, G.H., Wong-Riley, M., 1986. Laminar and cellular localization of cytochrome oxidase in the cat striate cortex. J. Comp. Neurol. 245, 137–159. 10.1002/cne.902450202

47. Kageyama, G.H., Wong-Riley, M.T., 1984. The histochemical localization of cytochrome oxidase in the retina and lateral geniculate nucleus of the ferret, cat, and monkey, with particular reference to retinal mosaics and ON/OFF-center visual channels. J. Neurosci. 4, 2445–2459. 10.1523/jneurosci.04-10-02445.1984

48. Kim, S.-G., Harel, N., Jin, T., Kim, T., Lee, P., Zhao, F., 2013. Cerebral blood volume MRI with intravascular superparamagnetic iron oxide nanoparticles. NMR Biomed. 26, 949–962. 10.1002/nbm.2885

49. Kimura, I., Hayashi, M.J., Amano, K., 2024. Immediate effect of quadri-pulse stimulation on human brain microstructures and functions. Imaging Neuroscience 2, 1–15. 10.1162/imag_a_00264

50. Kirilina, E., Helbling, S., Morawski, M., Pine, K., Reimann, K., Jankuhn, S., Dinse, J., Deistung, A., Reichenbach, J.R., Trampel, R., Geyer, S., Müller, L., Jakubowski, N., Arendt, T., Bazin, P.-L., Weiskopf, N., 2020. Superficial white matter imaging: Contrast mechanisms and whole-brain in vivo mapping. Sci Adv 6. 10.1126/sciadv.aaz9281

51. Lamsa, K.P., Kullmann, D.M., Woodin, M.A., 2010. Spike-timing dependent plasticity in inhibitory circuits. Front. Synaptic Neurosci. 2, 8. 10.3389/fnsyn.2010.00008

52. Lazari, A., Salvan, P., Cottaar, M., Papp, D., Rushworth, M.F.S., Johansen-Berg, H., 2022. Hebbian activity-dependent plasticity in white matter. Cell Rep. 39, 110951. 10.1016/j.celrep.2022.110951

53. Lee, K.H., Chung, K., Chung, J.M., Coggeshall, R.E., 1986. Correlation of cell body size, axon size, and signal conduction velocity for individually labelled dorsal root ganglion cells in the cat. J. Comp. Neurol. 243, 335–346. 10.1002/cne.902430305

54. Markov, N.T., Ercsey-Ravasz, M., Lamy, C., Ribeiro Gomes, A.R., Magrou, L., Misery, P., Giroud, P., Barone, P., Dehay, C., Toroczkai, Z., Knoblauch, K., Van Essen, D.C., Kennedy, H., 2013. The role of long-range connections on the specificity of the macaque interareal cortical network. Proc. Natl. Acad. Sci. U. S. A. 110, 5187–5192. 10.1073/pnas.1218972110

55. Markov, N.T., Ercsey-Ravasz, M.M., Ribeiro Gomes, A.R., Lamy, C., Magrou, L., Vezoli, J., Misery, P., Falchier, A., Quilodran, R., Gariel, M.A., Sallet, J., Gamanut, R., Huissoud, C., Clavagnier, S., Giroud, P., Sappey-Marinier, D., Barone, P., Dehay, C., Toroczkai, Z., Knoblauch, K., Van Essen, D.C., Kennedy, H., 2014. A weighted and directed interareal connectivity matrix for macaque cerebral cortex. Cereb. Cortex 24, 17–36. 10.1093/cercor/bhs270

56. Murray, C.D., 1926. The physiological principle of minimum work: II. Oxygen exchange in capillaries. Proc. Natl. Acad. Sci. U. S. A. 12, 299–304. 10.1073/pnas.12.5.299

57. Nashmi, R., Fehlings, M.G., 2001. Changes in axonal physiology and morphology after chronic compressive injury of the rat thoracic spinal cord. Neuroscience 104, 235–251. 10.1016/s0306-4522(01)00009-4

58. Ogawa, S., Menon, R.S., Tank, D.W., Kim, S.G., Merkle, H., Ellermann, J.M., Ugurbil, K., 1993. Functional brain mapping by blood oxygenation level-dependent contrast magnetic resonance imaging. A comparison of signal characteristics with a biophysical model. Biophys. J. 64, 803–812. 10.1016/S0006-3495(93)81441-3

59. Perge, J.A., Niven, J.E., Mugnaini, E., Balasubramanian, V., Sterling, P., 2012. Why do axons differ in caliber? J. Neurosci. 32, 626–638. 10.1523/JNEUROSCI.4254-11.2012

60. Razavi, M.S., Shirani, E., Kassab, G.S., 2018. Scaling laws of flow rate, vessel blood volume, lengths, and transit times with number of capillaries. Front. Physiol. 9. 10.3389/fphys.2018.00581

61. Saab, A.S., Tzvetanova, I.D., Nave, K.-A., 2013. The role of myelin and oligodendrocytes in axonal energy metabolism. Curr. Opin. Neurobiol. 23, 1065–1072. 10.1016/j.conb.2013.09.008

62. Sampaio-Baptista, C., Johansen-Berg, H., 2017. White Matter Plasticity in the Adult Brain. Neuron 96, 1239–1251. 10.1016/j.neuron.2017.11.026

63. Sampaio-Baptista, C., Neyedli, H.F., Sanders, Z.-B., Diosi, K., Havard, D., Huang, Y., Andersson, J.L.R., Lühr, M., Goebel, R., Johansen-Berg, H., 2021. fMRI neurofeedback in the motor system elicits bidirectional changes in activity and in white matter structure in the adult human brain. Cell Rep. 37, 109890. 10.1016/j.celrep.2021.109890

64. Sandrone, S., Aiello, M., Cavaliere, C., Thiebaut de Schotten, M., Reimann, K., Troakes, C., Bodi, I., Lacerda, L., Monti, S., Murphy, D., Geyer, S., Catani, M., Dell’Acqua, F., 2023. Mapping myelin in white matter with T1-weighted/T2-weighted maps: discrepancy with histology and other myelin MRI measures. Brain Struct. Funct. 228, 525–535. 10.1007/s00429-022-02600-z

65. Schilling, K.G., Li, M., Rheault, F., Gao, Y., Cai, L., Zhao, Y., Xu, L., Ding, Z., Anderson, A.W., Landman, B.A., Gore, J.C., 2023. Whole-brain, gray, and white matter time-locked functional signal changes with simple tasks and model-free analysis. Proc. Natl. Acad. Sci. U. S. A. 120, e2219666120. 10.1073/pnas.2219666120

66. Schilling, K.G., Newton, A., Tax, C.M.W., Chamberland, M., Remedios, S.W., Gao, Y., Li, M., Chang, C., Rheault, F., Sepherband, F., Anderson, A., Gore, J.C., Landman, B., 2025. The relationship of white matter tract orientation to vascular geometry in the human brain. Sci. Rep. 15, 18396. 10.1038/s41598-025-99724-z

67. Shu, Y., Hasenstaub, A., McCormick, D.A., 2003. Turning on and off recurrent balanced cortical activity. Nature 423, 288–293. 10.1038/nature01616

68. Smirnov, M., Destrieux, C., Maldonado, I.L., 2021. Cerebral white matter vasculature: still uncharted? Brain 144, 3561–3575. 10.1093/brain/awab273

69. Tabelow, K., Balteau, E., Ashburner, J., Callaghan, M.F., Draganski, B., Helms, G., Kherif, F., Leutritz, T., Lutti, A., Phillips, C., Reimer, E., Ruthotto, L., Seif, M., Weiskopf, N., Ziegler, G., Mohammadi, S., 2019. hMRI - A toolbox for quantitative MRI in neuroscience and clinical research. Neuroimage 194, 191–210. 10.1016/j.neuroimage.2019.01.029

70. Tahedl, M., Tournier, J.-D., Smith, R.E., 2025. Structural connectome construction using constrained spherical deconvolution in multi-shell diffusion-weighted magnetic resonance imaging. Nat. Protoc. 10.1038/s41596-024-01129-1

71. Tao, K., Matsuki, N., Koyama, R., 2014. AMP-activated protein kinase mediates activity-dependent axon branching by recruiting mitochondria to axon: Role of Mitochondria in Axonal Morphogenesis. Dev. Neurobiol. 74, 557–573. 10.1002/dneu.22149

72. Tournier, J.-D., Smith, R., Raffelt, D., Tabbara, R., Dhollander, T., Pietsch, M., Christiaens, D., Jeurissen, B., Yeh, C.-H., Connelly, A., 2019. MRtrix3: A fast, flexible and open software framework for medical image processing and visualisation. Neuroimage 202, 116137. 10.1016/j.neuroimage.2019.116137

73. Van Essen, D.C., Maunsell, J.H., 1980. Two-dimensional maps of the cerebral cortex. J. Comp. Neurol. 191, 255–281. 10.1002/cne.901910208

74. Vezoli, J., Vinck, M., Bosman, C.A., Bastos, A.M., Lewis, C.M., Kennedy, H., Fries, P., 2021. Brain rhythms define distinct interaction networks with differential dependence on anatomy. Neuron 109, 3862–3878.e5. 10.1016/j.neuron.2021.09.052

75. Viessmann, O., Tian, Q., Bernier, M., Polimeni, J.R., 2022. Static and dynamic BOLD fMRI components along white matter fibre tracts and their dependence on the orientation of the local diffusion tensor axis relative to the B0-field. J. Cereb. Blood Flow Metab. 42, 1905–1919. 10.1177/0271678X221106277

76. Wang, H., Wang, X., Wang, Y., Zhang, D., Yang, Y., Zhou, Y., Qiu, B., Zhang, P., 2023. White matter BOLD signals at 7 Tesla reveal visual field maps in optic radiation and vertical occipital fasciculus. Neuroimage 269, 119916. 10.1016/j.neuroimage.2023.119916

77. Weber, B., 2002. White matter glucose metabolism during intracortical electrostimulation: A quantitative [18F]fluorodeoxyglucose autoradiography study in the rat. Neuroimage 16, 993–998. 10.1006/nimg.2002.1104

78. Weber, B., Keller, A.L., Reichold, J., Logothetis, N.K., 2008. The microvascular system of the striate and extrastriate visual cortex of the macaque. Cereb. Cortex 18, 2318–2330. 10.1093/cercor/bhm259

79. Wiser, A.K., Callaway, E.M., 1996. Contributions of individual layer 6 pyramidal neurons to local circuitry in macaque primary visual cortex. J. Neurosci. 16, 2724–2739. 10.1523/jneurosci.16-08-02724.1996

80. Wong-Riley, M.T., 1989. Cytochrome oxidase: an endogenous metabolic marker for neuronal activity. Trends Neurosci. 12, 94–101. 10.1016/0166-2236(89)90165-3

81. Wu, W.-C., Lin, S.-C., Wang, D.J., Chen, K.-L., Li, Y.-D., 2013. Measurement of cerebral white matter perfusion using pseudocontinuous arterial spin labeling 3T magnetic resonance imaging--an experimental and theoretical investigation of feasibility. PLoS One 8, e82679. 10.1371/journal.pone.0082679

82. Yang, S., Park, J.H., Lu, H.-C., 2023. Axonal energy metabolism, and the effects in aging and neurodegenerative diseases. Mol. Neurodegener. 18, 49. 10.1186/s13024-023-00634-3

83. Yeatman, J.D., Dougherty, R.F., Myall, N.J., Wandell, B.A., Feldman, H.M., 2012. Tract profiles of white matter properties: automating fiber-tract quantification. PLoS One 7, e49790. 10.1371/journal.pone.0049790

84. Zhang, H., Schneider, T., Wheeler-Kingshott, C.A., Alexander, D.C., 2012. NODDI: Practical in vivo neurite orientation dispersion and density imaging of the human brain. Neuroimage 61, 1000–1016. 10.1016/j.neuroimage.2012.03.072

85. Zheng, D., LaMantia, A.S., Purves, D., 1991. Specialized vascularization of the primate visual cortex. J. Neurosci. 11, 2622–2629. 10.1523/jneurosci.11-08-02622.1991

