## Supplementary Information for "Linking neuron-axon-synapse architecture to white matter vasculature using high-resolution multimodal MRI in primate brain"

Contains Supplementary Figures 1–5 and Table 1

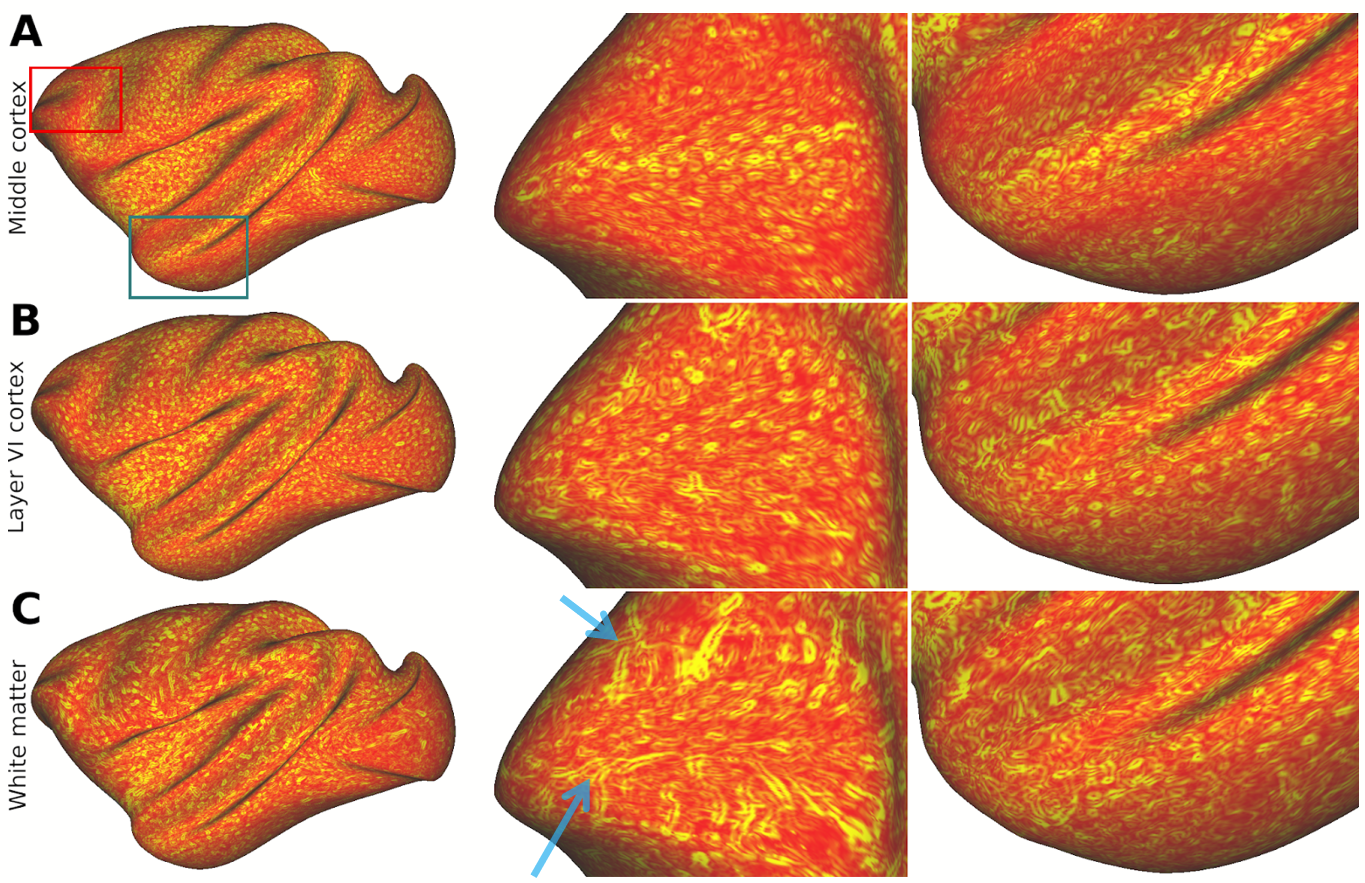

**Supplementary Figure 1.** **Regional variations in superficial white matter macrovasculature patterns across the brain.** Spatial gradients of the ferumoxytol-weighted image are shown for the equivolumetric layers (EL): (A) EL4a, located near the cortical midthickness, and (B) EL6b, situated adjacent to the white matter surface. (C) An equidistant superficial white matter surface located beneath the cortical gray matter. Distinct and variable vessel orientations were observed in the prefrontal cortex (red box; middle panels), whereas subtler variations in vessel orientation were evident near the temporal pole (blue box; right panels). Blue arrows indicate exemplar vessels oriented orthogonal to each other.

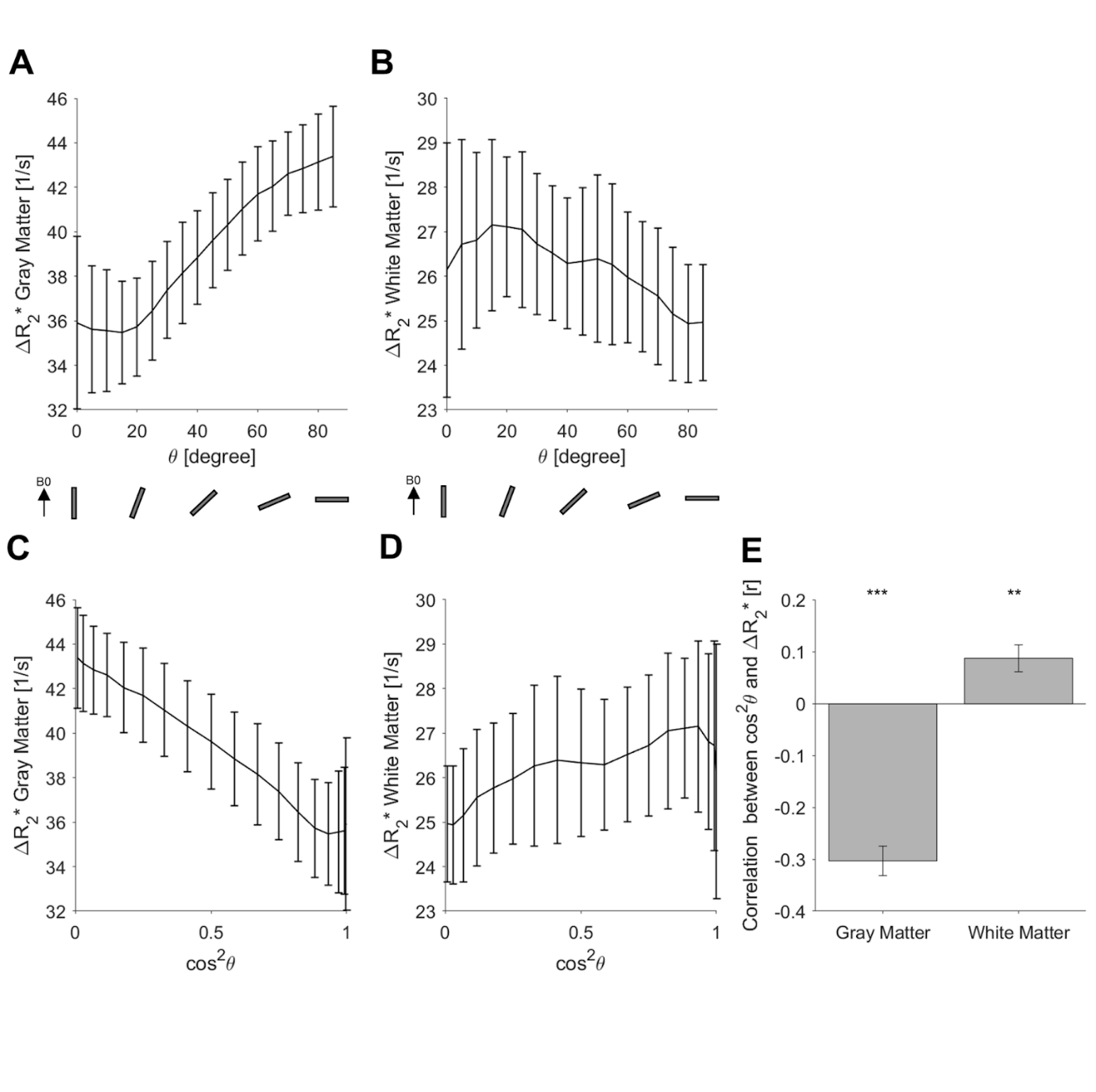

**Supplementary Figure 2. Relationship between ferumoxytol-induced change in transverse relaxation rate (ΔR_2_*) and the angle (θ) between the normal of cortex and B_0_. (A)** Relationship between ΔR_2_* and θ in the cortical gray matter. **(B)** Relationship between ΔR_2_* and θ in the superficial white matter. **(C)** Relationship between ΔR_2_* and ${cos}^{2}\theta$ in the cortical gray matter. **(D)** Relationship between ΔR_2_* and ${cos}^{2}\theta$ in the superficial white matter. **(E)** Pearson’s correlation coefficients between ΔR_2_* and the ${cos}^{2}\theta$ within the cortical gray matter (left) and superficial white matter (right). Linear relationship is more prominent in the cortical gray matter probably due to more systematic orientation of large vessels. ** *P* < 0.005 and ** *P* < 0.001 (Bonferroni-corrected).

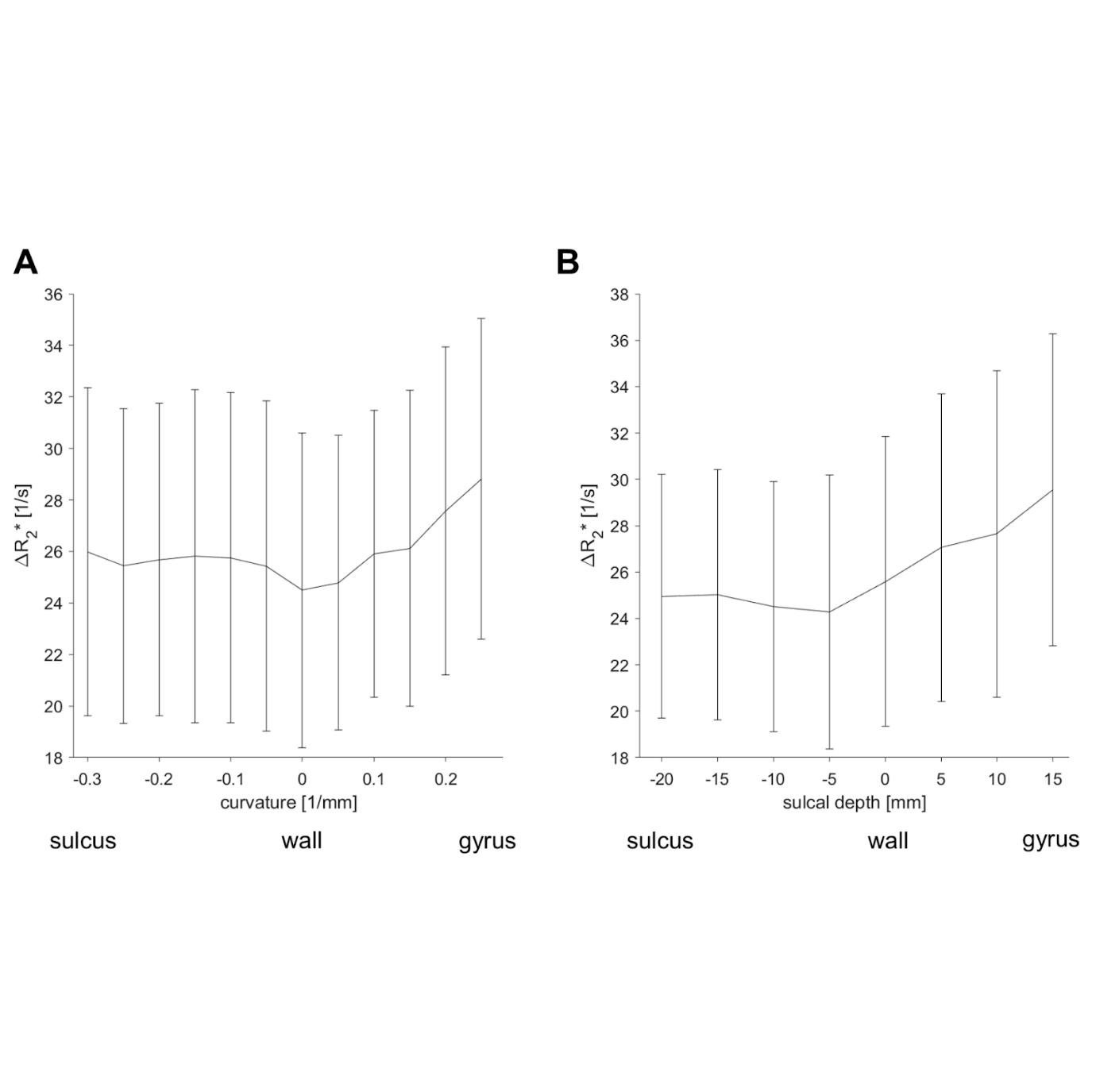

**Supplementary Figure 3. Relationship between superficial white matter vasculature and cortical geometry.** Ferumoxytol-induced change in transverse relaxation rate **(**ΔR_2_*) plotted as a function of **(A)** curvature and **(B)** sulcal depth. In both plots, negative values correspond to sulcus and positive values indicate gyrus.

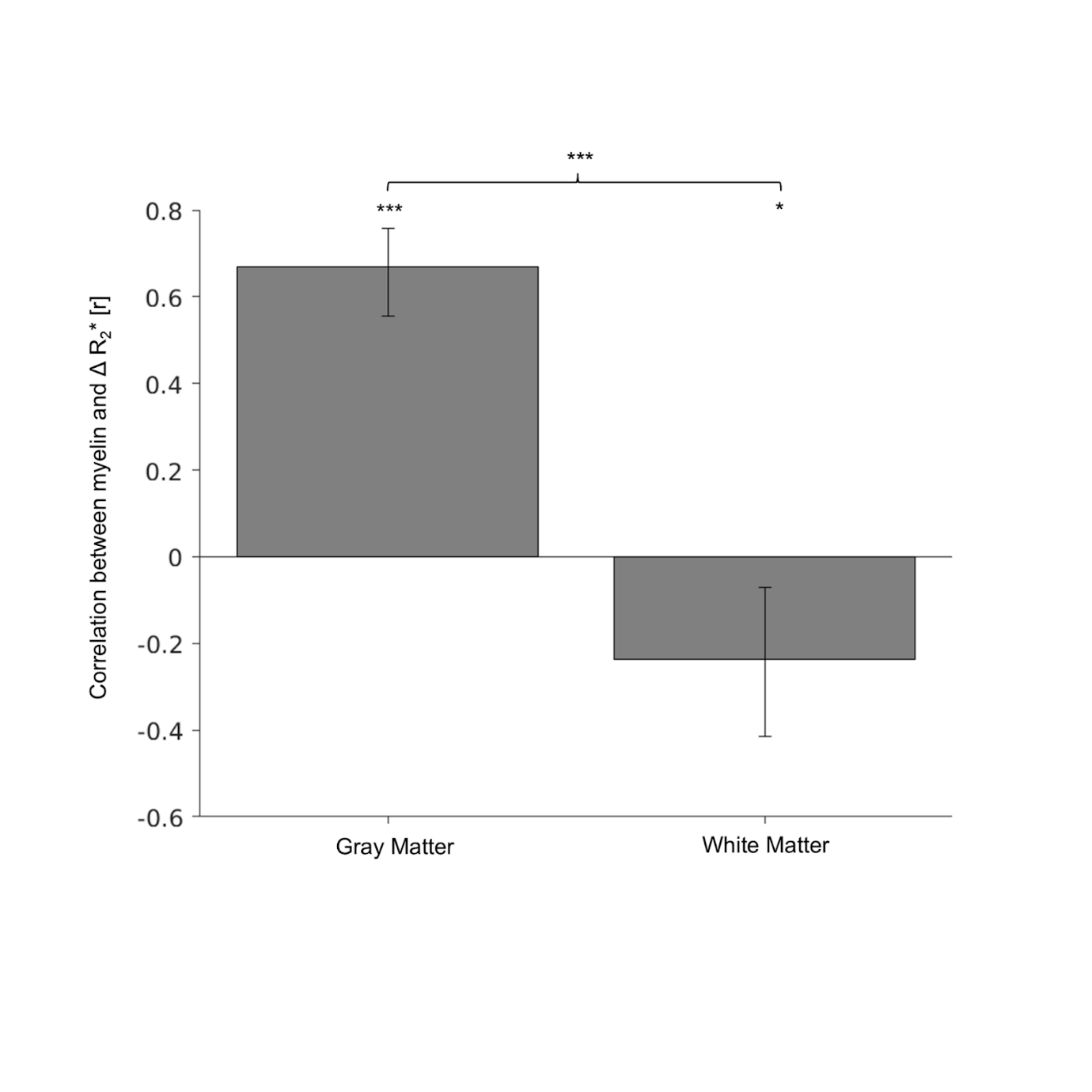

**Supplementary Figure 4. Contrasting relationship between blood volume and myelin acorss superficial white matter and cortical gray matter.** Correlation between ΔR_2_*, an indirect proxy measure of vascular volume, and T1w/T2w-FLAIR, an indirect proxy measure of myelin density. Data were parcellated using the M132 macaque atlas (See Methods). ** *P* < 0.005 and ** *P* < 0.001 (Bonferroni-corrected).

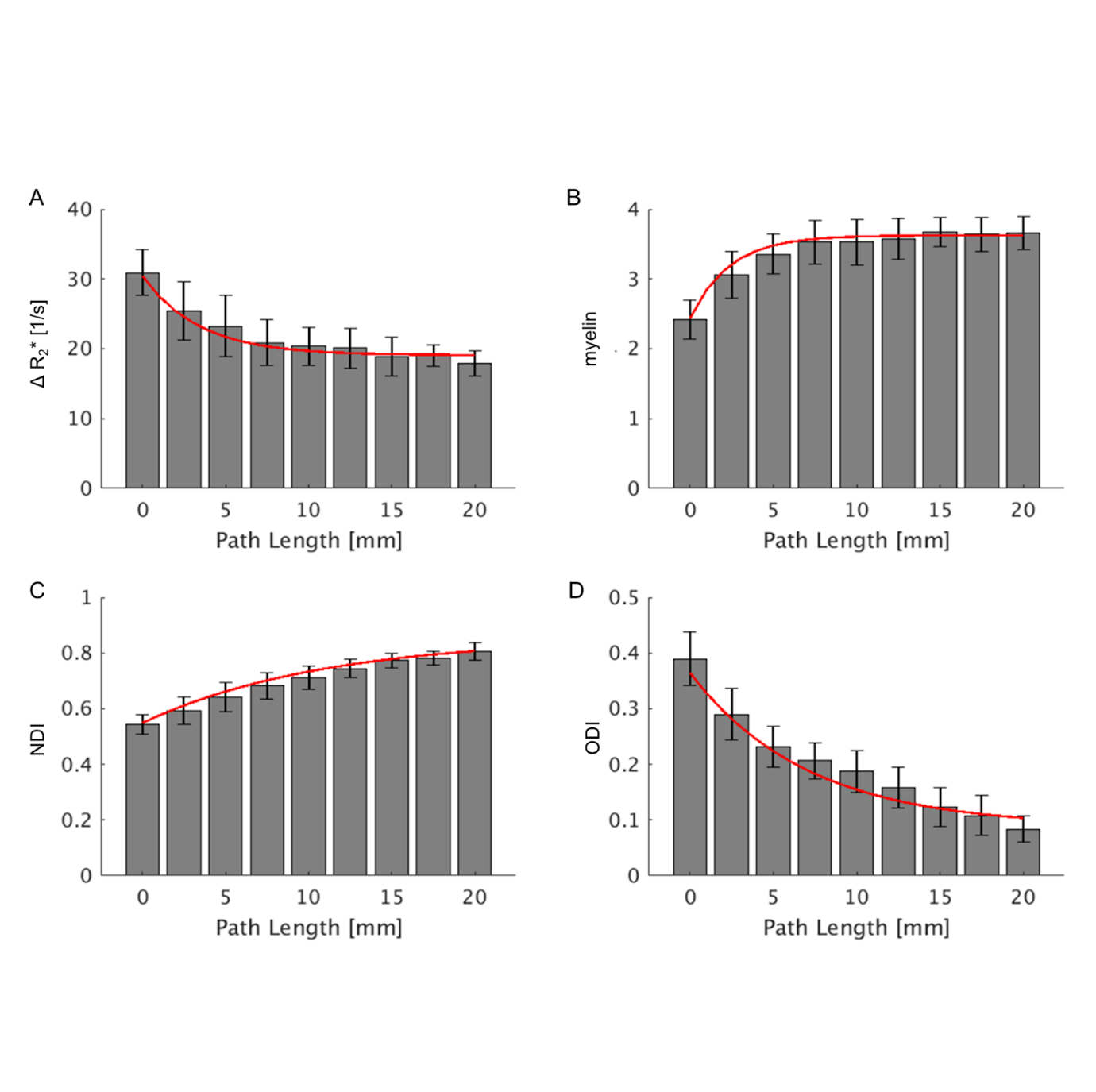

**Supplementary Figure 5. Relationships between white matter vascularity, microstructure, and brain geometry.** The relationships between path length from superficial white matter and **(A)** ferumoxytol-induced change in transverse relaxation rate (ΔR_2_*), **(B)** T1w/T2w-FLAIR, an indirect proxy measure of myelin density, **(C)** neurite density index (NDI), and **(D)** orientation dispersion index (ODI).

|  | Exponential | Linear | Quadratic | Logistic |
| --- | --- | --- | --- | --- |
| ΔR_2_*    AICc    BIC  NDI    AICc    BIC  ODI    AICc    BIC | **1717**  **1728**  **-1086**  **-1075**  **-1172**  **-1161** | 1804  1811  -1061  -1053    -1112  -1104 | 1725  1736  -1085  -1073  -1156  -1145 | 1718  1733  -1085  -1070  -1171  -1156 |

**Table S1. Model comparisons for fitting each metric as a function of relative distance.** The bolded values indicate the lowest AICc and BIC scores across models, corresponding to the best-fitting model. ΔR_2_*: transverse relaxation rate; NDI: neurite density index; ODI: orientation dispersion index; AICc: corrected Akaike Information Criterion; BIC: Bayesian Information Criterion.
